# A ribosomal pseudouridine drives an oncogenic translation program

**DOI:** 10.64898/2026.10.06.757013

**Authors:** Laurence Faucher-Giguère, Andrea Rivera-Pizana, Baudouin Seguineau de Préval, Étienne Fafard-Couture, Gabrielle Deschamps-Francoeur, Virginie Marchand, Yuri Motorin, Charles Barchet, Bruno P. Klaholz, Taha Azad, Michelle S. Scott, Sherif Abou Elela

## Abstract

Ribosome heterogeneity can regulate translation, but how changes in ribosome chemistry reshape cellular programs and contribute to diseases such as cancer remains incompletely understood. Here we show that SNORA81-guided pseudouridylation at a single 28S ribosomal RNA site, U4636, links ribosome heterogeneity to selective translation and ovarian cancer cell proliferation. This partially modified site contacts eL31 near the peptide-exit region. SNORA81 depletion reduces U4636 pseudouridylation and changes ribosome occupancy in relation to codon identity and nascent-peptide properties without broadly disrupting rRNA processing or bulk polysome profiles. Depletion also redistributes mRNAs with distinct GC content and amino-acid composition across polysomes, with corresponding protein-abundance changes in selected cancer-associated targets. Wild-type SNORA81, but not a guide-region mutant, restores proliferation. Together, these findings support a “sno-balling” model in which altered abundance of a single snoRNA propagates through rRNA modification, selective ribosome engagement and protein output to reshape cancer-associated gene expression and cell behavior.

**Graphical Abstract:** The sno-balling effect links a single rRNA modification to cancer-associated translation. SNORA81-guided pseudouridylation at U4636 is associated with changes in ribosome occupancy, polysome association of compositionally distinct STAR mRNAs, and selected protein abundances. These findings support a model linking variation in rRNA modification to ovarian cancer cell proliferation.

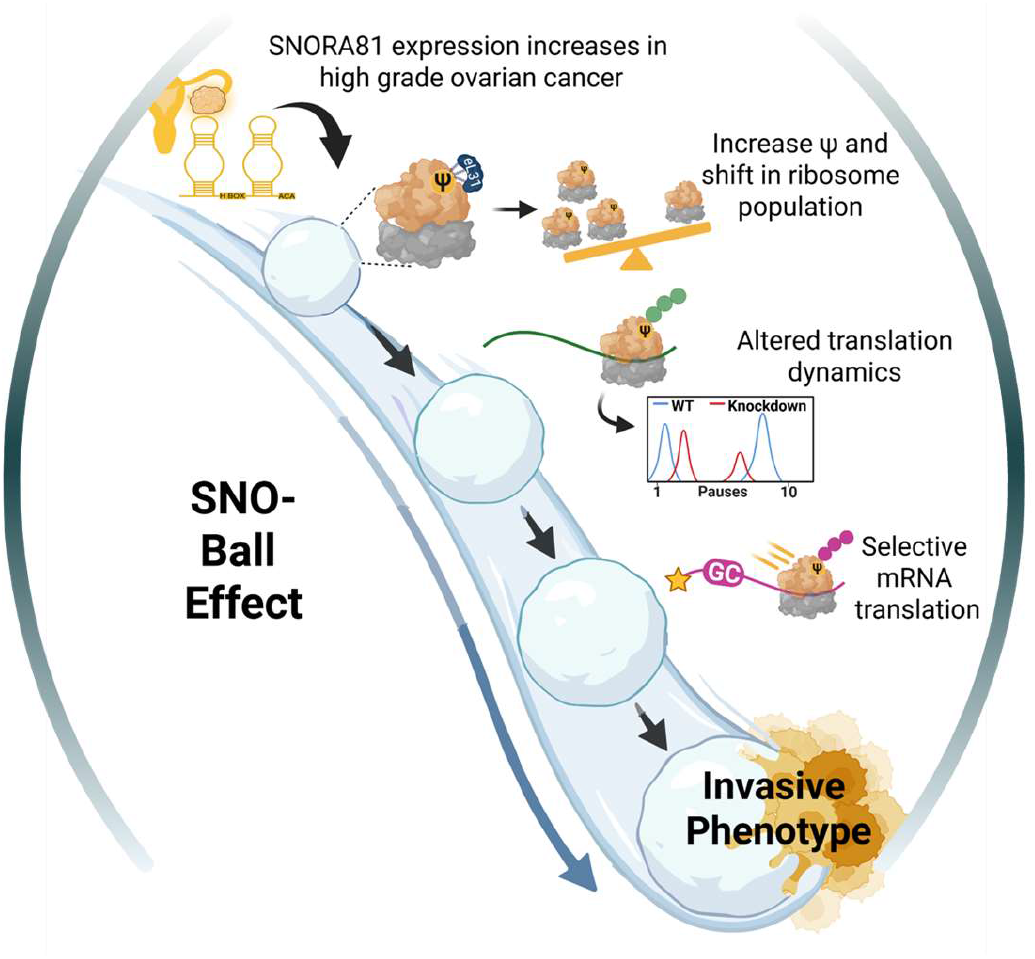

## INTRODUCTION

Ribosomes are increasingly recognized as heterogeneous machines whose composition and chemical state can shape translational output^1–3^. This heterogeneity arises from variation in ribosomal proteins, ribosomal RNA (rRNA) sequences and modifications, and associated factors^1–3^. rRNA modifications, including pseudouridylation and 2′-O-methylation, are a major source of this heterogeneity and can influence ribosome structure, decoding and dynamics^4–8^. Although many rRNA sites are constitutively modified, others are modified in only a fraction of ribosomes, and this fraction varies across tissues, developmental states and diseases^4,9^. How changes in ribosome chemistry reshape coordinated gene expression programs and contribute to cellular phenotypes remains incompletely understood.

Many rRNA modifications are directed by small nucleolar RNAs (snoRNAs), which recognize specific target sequences. Dysregulation of snoRNAs in cancer can therefore connect altered ribosome modification to changes in ribosome activity and tumour phenotypes^6,10^. For example, loss of the H/ACA snoRNA SNORA24, which guides pseudouridylation at two sites in 18S rRNA, alters ribosome decoding and translational fidelity and cooperates with oncogenic RAS to promote liver cancer^11^. Similarly, MYC signaling alters site-specific rRNA 2′-O-methylation, and removal of one such modification changes the translation of selected mRNAs and associated cellular phenotypes^12^. These studies establish rRNA modification as a regulator of ribosome function and cellular behavior. However, the relationships between individual modifications, the sequence features of responsive mRNAs and coordinated cancer-associated protein changes remain poorly defined.

To address these questions, we focused on SNORA81, an H/ACA snoRNA that is upregulated in high-grade serous ovarian carcinoma (HGSC) and whose abundance correlates with rRNA modification, proliferation and migration^13^. SNORA81 guides pseudouridylation of U4636 in helix 96 of 28S rRNA. U4636 lies on the outer surface of the 60S subunit, at an rRNA-eL31 interface near the peptide-exit region, far from the decoding and catalytic centers. We asked how SNORA81-dependent pseudouridylation at U4636 influences ribosome behavior, the engagement of compositionally distinct mRNAs and cancer cell proliferation. We find that SNORA81-guided pseudouridylation at U4636 (Ψ4636) supports ovarian cancer cell proliferation. SNORA81 depletion also alters ribosome occupancy associated with codon identity and nascent-peptide properties, redistributes compositionally distinct mRNAs across polysomes and changes the abundance of selected cancer-associated proteins. Together, these findings link variation at a single rRNA modification site to sequence-associated changes in ribosome behavior, selective mRNA engagement and cancer-associated protein expression.

## RESULTS

### Ψ4636 lies at an rRNA-eL31 interface

SNORA81 was previously identified as part of an H/ACA snoRNA signature that distinguishes HGSC from serous borderline tumours (SBT) and normal ovarian tissue^13^. Its abundance correlates with pseudouridylation at its guided 28S rRNA target site, and SNORA81 knockdown reduces pseudouridylation at this site and suppresses ovarian cancer cell proliferation and migration, establishing SNORA81 as a candidate driver of disease progression (**Figure 1A**).

**Figure 1:**
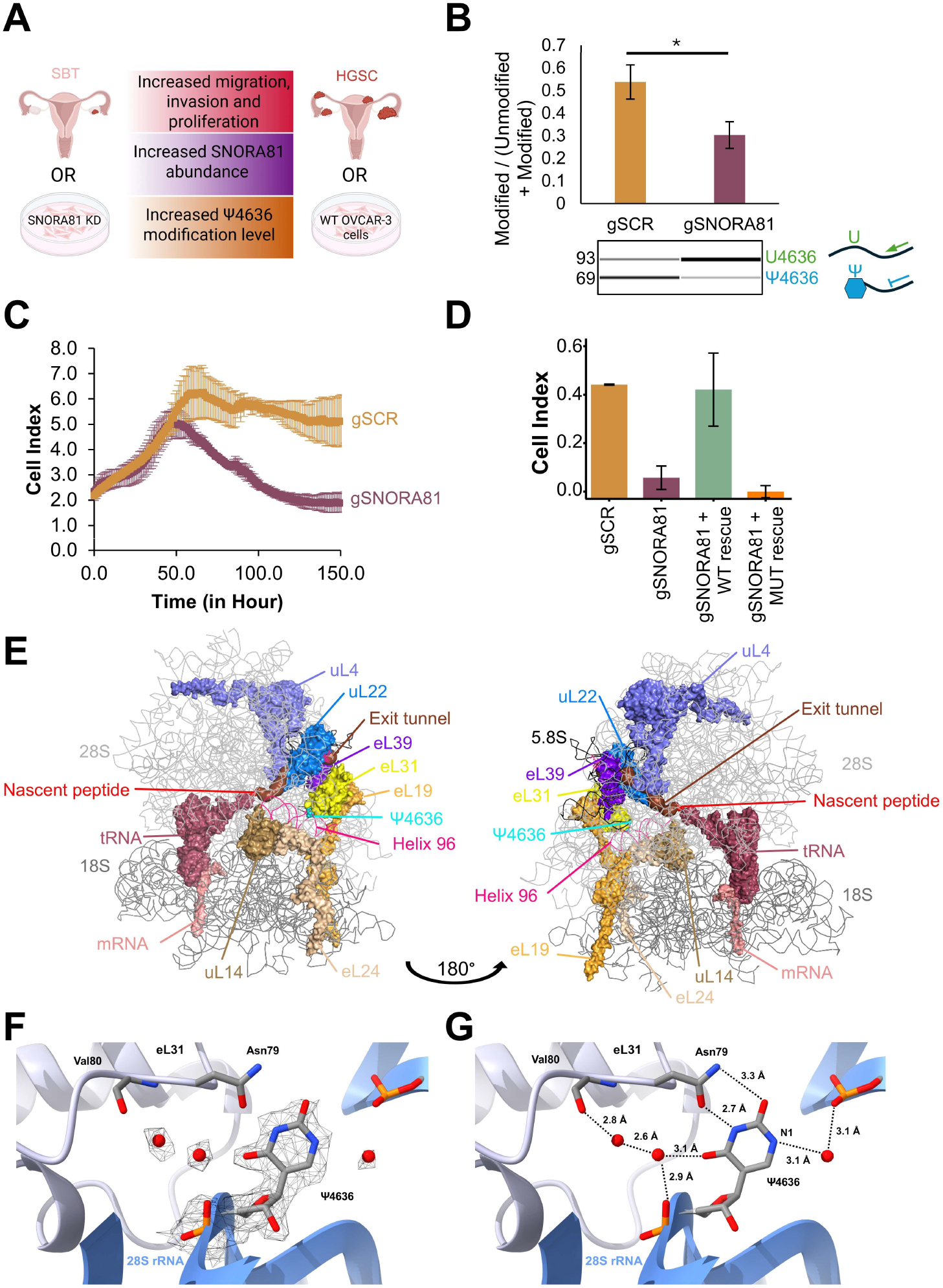
SNORA81 directs pseudouridylation at U4636 near the peptide exit tunnel. **(A)** Schematic integrating observations from ovarian tumour samples and OVCAR-3 cells. Serous borderline tumour (SBT) samples and SNORA81-depleted cells have lower SNORA81 abundance and Ψ4636 modification level, whereas HGSC samples and wild-type OVCAR-3 cells have higher levels. Higher SNORA81 abundance and Ψ4636 modification level are associated with increased migration, invasion and proliferation. Created with BioRender. **(B)** Quantification of U4636 pseudouridylation by CLAP after CRISPR–Cas13d-mediated SNORA81 depletion (gSNORA81) compared with scrambled control (gSCR). Values indicate the modified fraction. The schematic shows the 93-nt read-through PCR product from unmodified U4636 and the 69-nt RT-stop product generated by CMC-modified Ψ4636. n = 3 biologically independent samples; two-sided t-test assuming unequal variance p = 0.015; bars, mean ± s.d. **(C)** xCELLigence analysis of OVCAR-3 cells after SNORA81 depletion. Cell Index, an impedance-based readout of adherent cell number, morphology and viability, was reduced in gSNORA81 cells. Data are representative of three independent experiments; points, mean of two technical replicates; error bars, s.d. **(D)** xCELLigence analysis at 150 hours after rescue with wild-type (WT) or guide-region mutant SNORA81. Data are representative of three independent experiments; bars, mean of two technical replicates; error bars, s.d. **(E)** Views of the human ribosome structure^18^ showing Ψ4636 in 28S rRNA near eL31 and peptide-exit-tunnel proteins. **(F)** Local environment of Ψ4636 (PDB 8QOI) with the experimental density of the cryo-EM map. **(G)** Ψ4636 forms reciprocal hydrogen bonds with Asn79 of eL31 and participates in a water-mediated hydrogen-bond network involving the backbone of eL31 residue Val80.

To assess tumour-type differences in pseudouridylation at individual rRNA sites, we directly quantified rRNA pseudouridylation across HGSC and SBT samples by HydraPsiSeq^14^, which enables parallel quantification of pseudouridylation at multiple rRNA sites. Eighteen sites showed an absolute difference in mean modified fraction greater than 0.05 between tumour types, with five sites more highly pseudouridylated in HGSC and thirteen more highly pseudouridylated in SBT (**Figure S1A, B**). U4636 was among the five sites with higher mean pseudouridylation in HGSC, and its guide snoRNA, SNORA81, was one of two guide snoRNAs that were also significantly more abundant in HGSC in previously published RNA-sequencing data^13^ (**Figure S1C**). The other, SNORA56, was also part of the HGSC-associated snoRNA signature identified previously^13^. Together, these comparisons prioritized SNORA81-guided pseudouridylation at U4636 for functional investigation.

To examine the consequences of reducing SNORA81-guided pseudouridylation at U4636, we depleted SNORA81 with CRISPR–Cas13d without altering the abundance of its host gene, *EIF4A2* (**Figure S2A, B**). In OVCAR-3 cells, a model epithelial ovarian cancer cell line, CMC–RT and ligation-assisted PCR (CLAP^15^) confirmed reduced pseudouridylation at U4636 following SNORA81 depletion (**Figure 1B**). Long-read nanopore sequencing 48 hours after depletion was used to determine whether depletion also altered pseudouridylation at other rRNA sites. We detected no consistent changes at other pseudouridylation sites above the detection threshold of the assay, indicating that SNORA81 depletion reduces pseudouridylation at U4636 without detectably remodelling the broader rRNA pseudouridylation landscape (**Figure S2C**). SNORA81 depletion also reduced the OVCAR-3 cell index, consistent with impaired proliferation (**Figure 1C**). To determine whether the reduced proliferation reflected a general defect in ribosome production, we next assessed ribosome biogenesis and integrity. Ribosomal-protein mRNA abundance was broadly unchanged by RNA-seq (**Figure S3A**), the 28S/18S rRNA ratio measured from equal cell numbers remained similar across the examined time points (**Figure S3B**), and pre-rRNA processing intermediates showed no major defects (**Figure S3C**). Together, these measurements provided no evidence of a broad defect in ribosome biogenesis or pre-rRNA processing following SNORA81 depletion. Re-expression of wild-type SNORA81 restored proliferation, whereas a guide-region mutant designed to disrupt U4636 targeting while preserving predicted snoRNA structure and expression, based on snoReport^16^ and a snoRNA expression predictor^17^, failed to rescue growth (**Figure 1D**). Together with the measured reduction in pseudouridylation at U4636, these rescue results support a modification-dependent contribution of U4636 to the SNORA81 growth phenotype.

To investigate how U4636 might affect cell growth, we examined its position within the ribosome. Mapping these variable pseudouridylation sites onto the human 80S ribosome showed that many variable sites localized to the periphery of the large subunit rather than to the catalytic core, with a subset near the peptide exit tunnel (**Figure S1D**). U4636 exemplifies this pattern: it lies in helix 96 (H96) of 28S rRNA on the outer surface of the 60S subunit, at an rRNA–protein interface with eL31 near the peptide-exit region (**Figure 1E**). In the human 80S structure, eL31 forms part of the ribosomal surface surrounding the peptide-exit region^18^.

Available human ribosome structures show a largely conserved molecular environment around the U4636 position, with some variation in the conformation of the base and its interactions with eL31 (**Figure S4A**). H96 forms a kissing-loop interaction with helix 57 (H57), contributing to the organization of this rRNA region (**Figure S4B**). U4636 is not universally conserved across eukaryotes but is retained in most of the mammalian sequences examined and in several representatives of fungi and protists, with greater variation among more distantly related lineages (**Figure S4C**). H96 forms part of the H94–H97 four-way junction, placing U4636 within a broader rRNA structural framework (**Figure S4D**). To examine the structural environment of Ψ4636 in detail, we analysed the 1.9-Å human ribosome structure containing this modification (PDB 8QOI)^18^. The cryo-EM density supports the assignment of Ψ4636, including density assigned to a water molecule adjacent to N1, and the modification was independently validated by mass spectrometry^18^ (**Figure 1F**). Ψ4636 is flipped out of H96 and does not form a Watson–Crick base pair with another nucleotide; instead, it forms reciprocal hydrogen bonds with the side-chain amide of Asn79 in eL31 (**Figure 1G**). Ψ4636 adopts a defined orientation at the rRNA–eL31 interface and participates in a hydrogen-bond network involving two water molecules and the backbone of eL31 residue Val80 (**Figure 1G**). Thus, Ψ4636 forms a defined contact with eL31 near the peptide-exit region, providing a possible structural basis for its functional effects.

### Ψ4636 shapes specific codon and peptide pausing

We next asked whether U4636 pseudouridylation influences translation dynamics in OVCAR-3 cells, a model of high-grade serous ovarian cancer. To capture early translational changes while minimizing secondary effects, we performed ribosome profiling 24 h after SNORA81 depletion, before measurable proliferation or viability defects emerged (**Figure 2A**). Ribosome-profiling libraries passed quality control and showed comparable coding-sequence enrichment, transcript coverage and reading-frame assignment across conditions (**Figure 2B; Figure S5**). Metagene analysis around annotated start and stop codons yielded different inferred P-site offsets between conditions (-9 nt for gSCR and -12 nt for gSNORA81), although the overall profiles remained similar (**Figure 2C**). The 3-nt difference was reproducible across biological replicates and retained read-length classes. However, because P-site offsets are inferred from ribosome-protected fragment boundaries, this observation cannot distinguish altered ribosome positioning or protection geometry from systematic differences in footprint generation or processing. Although not mechanistically diagnostic, its reproducibility prompted us to ask whether it was accompanied by changes in codon-level P-site occupancy.

**Figure 2:**
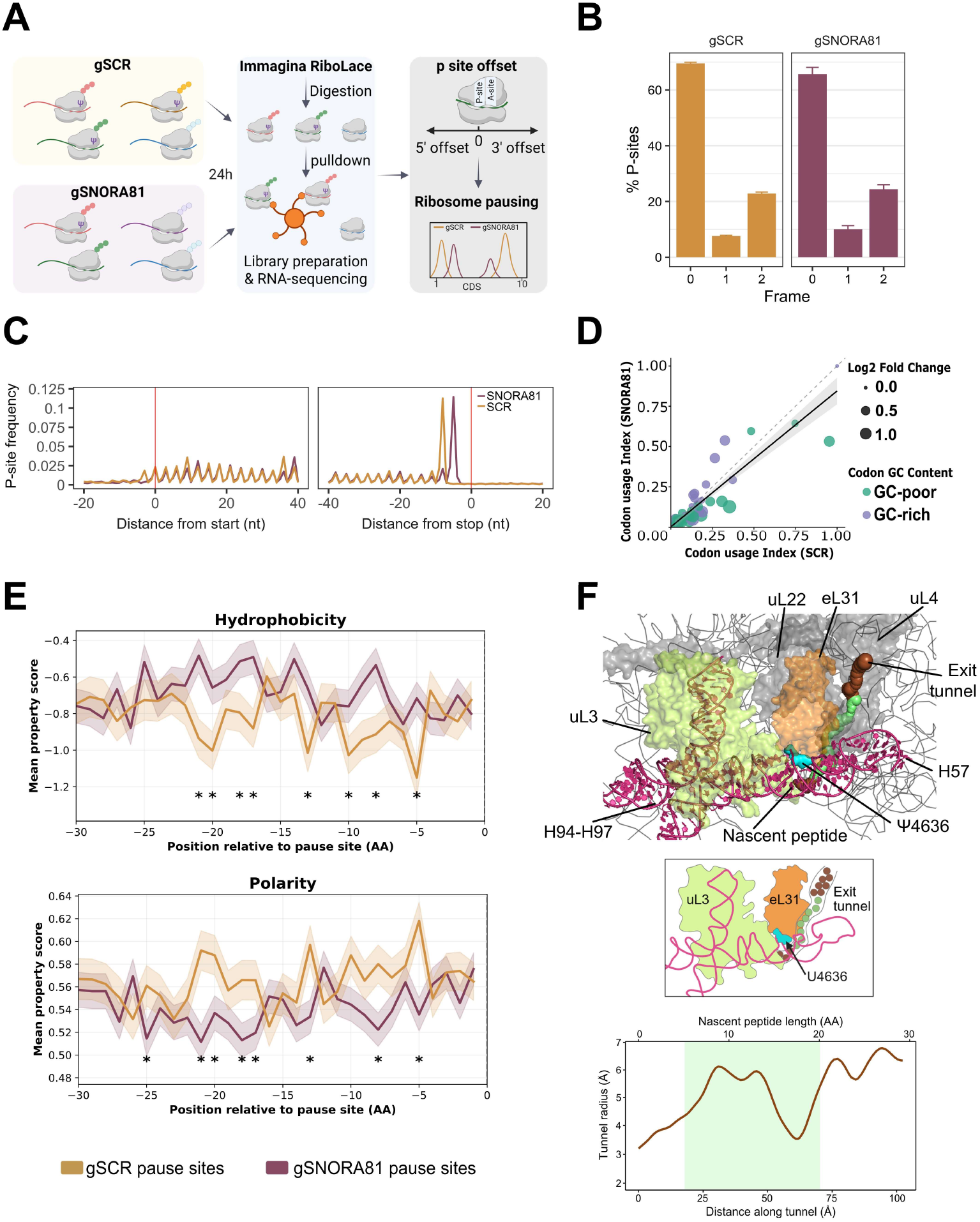
SNORA81-guided Ψ4636 remodels codon- and nascent-peptide-associated pausing. **(A)** Ribosome-profiling workflow used to quantify ribosome occupancy and P-site pausing 24 hours after CRISPR–Cas13d-mediated SNORA81 depletion in OVCAR-3 cells. Ribosome-protected fragments from gSCR (control guide) and gSNORA81 (SNORA81-targeting guides) cells were mapped to coding sequences and assigned to ribosomal P sites. Created with BioRender. **(B)** Distribution of mapped P sites across the three coding frames in gSCR and gSNORA81 samples. Data are mean ± s.d. from n = 4 biologically independent samples. **(C)** Metagene profiles of normalized P-site density around annotated start codons and stop codons. **(D)** Codon-level comparison of P-site occupancy in gSCR and gSNORA81 cells. Each point represents one codon; colour denotes GC content and point size indicates the absolute log2 fold change in pausing after SNORA81 depletion. Solid line, linear regression; dashed line, identity line. **(E)** Hydrophobicity and polarity of nascent-peptide windows 0-30 amino acids upstream of gSCR and gSNORA81 enriched pause sites. Significance at each position was assessed using two-sided Mann–Whitney U tests; P < 0.05. **(F)** Structural mapping of the SNORA81-sensitive peptide segment onto the ribosome exit tunnel. The segment corresponding to positions −5 to −20 upstream of pause sites is highlighted in green. The top panel shows the exit tunnel and uL4/uL22 constriction and uL3 interaction; the bottom panel shows the corresponding tunnel-radius profile.

At codon resolution, SNORA81 depletion selectively altered P-site occupancy (**Figure 2D**). GC-rich codons showed higher occupancy after depletion, whereas AU-rich codons showed relatively higher occupancy in control cells. Because Ψ4636 lies near the peptide exit region, where nascent-chain interactions can influence elongation, we examined the amino-acid context surrounding differential pause sites. We identified 1,192 pause sites enriched upon SNORA81 depletion and 898 enriched in control cells. Peptide windows upstream of depletion-enriched pause sites showed distinct physicochemical properties, including greater hydrophobicity and lower polarity than control-enriched sites (**Figure 2E**). Consistent with this pattern, *KDELR1* and *STARD7* showed pronounced SNORA81-dependent pausing in hydrophobic regions, while *CD164* and *ATP5F1B* showed control-enriched pausing in polar regions (**Figure S6**). Together, these results link reduced Ψ4636 to codon- and peptide-associated pausing.

To relate these sequence features to ribosome structure, we estimated where the corresponding nascent-peptide segment would lie along the peptide exit tunnel. Assuming a fully extended nascent chain (approximately 3.5 Å per residue), residues 5–20 upstream of the pause site would extend from the PTC-proximal portion of the tunnel towards and through the constriction formed by uL4 and uL22 (formerly L17) (**Figure 2F**). The H96 internal loop containing Ψ4636 forms part of the H94–H97 four-way junction, which is positioned adjacent to eL31 and uL3. uL3 extends from this region towards the PTC and the uL4/uL22 constriction and has been implicated in coordinating functional transitions during translation elongation^19,20^. This organization places Ψ4636 within a structural region connected to the portion of the exit tunnel predicted to contain residues upstream of the pause site.

This structural placement is relevant because nascent-peptide properties can shape ribosome pausing. Hydrophobicity, charge and volume influence interactions between the nascent peptide and the exit tunnel^21–23^. The residues showing the strongest SNORA81-dependent differences in physicochemical properties are predicted to occupy this region of the tunnel, providing a potential link between reduced Ψ4636 and altered elongation. These findings support a model in which a single rRNA pseudouridine couples local ribosome architecture to elongation dynamics determined by codon identity and nascent-peptide properties.

### Ψ4636 redistributes mRNAs on polysomes

We next asked whether the pausing changes observed after SNORA81 depletion were accompanied by altered ribosome engagement of specific mRNAs. To test this, we performed polysome profiling coupled with fraction-specific RNA-seq to quantify transcript-level ribosome association (**Figure 3A**). We measured ribosome association as polysome enrichment relative to input and defined transcripts with extreme changes in polysome association (|z-score| ≥ 2) as SNORA81-dependent translationally altered RNAs (STAR mRNAs; **Figure 3A; Figure S7A**). Here, “translationally altered” refers specifically to altered ribosome association measured by polysome profiling. We refer to transcripts with increased ribosome association after SNORA81 depletion as over-associated and those with decreased association as under-associated. Bulk polysome profiles and the overall distribution of detected mRNAs across fractions were broadly similar after SNORA81 depletion (**Figure 3B; Figure S7B**). Despite this, 524 mRNAs met the STAR classification thresholds, comprising 289 under-associated and 235 over-associated transcripts (**Figure 3C; Figure S7A**).

**Figure 3:**
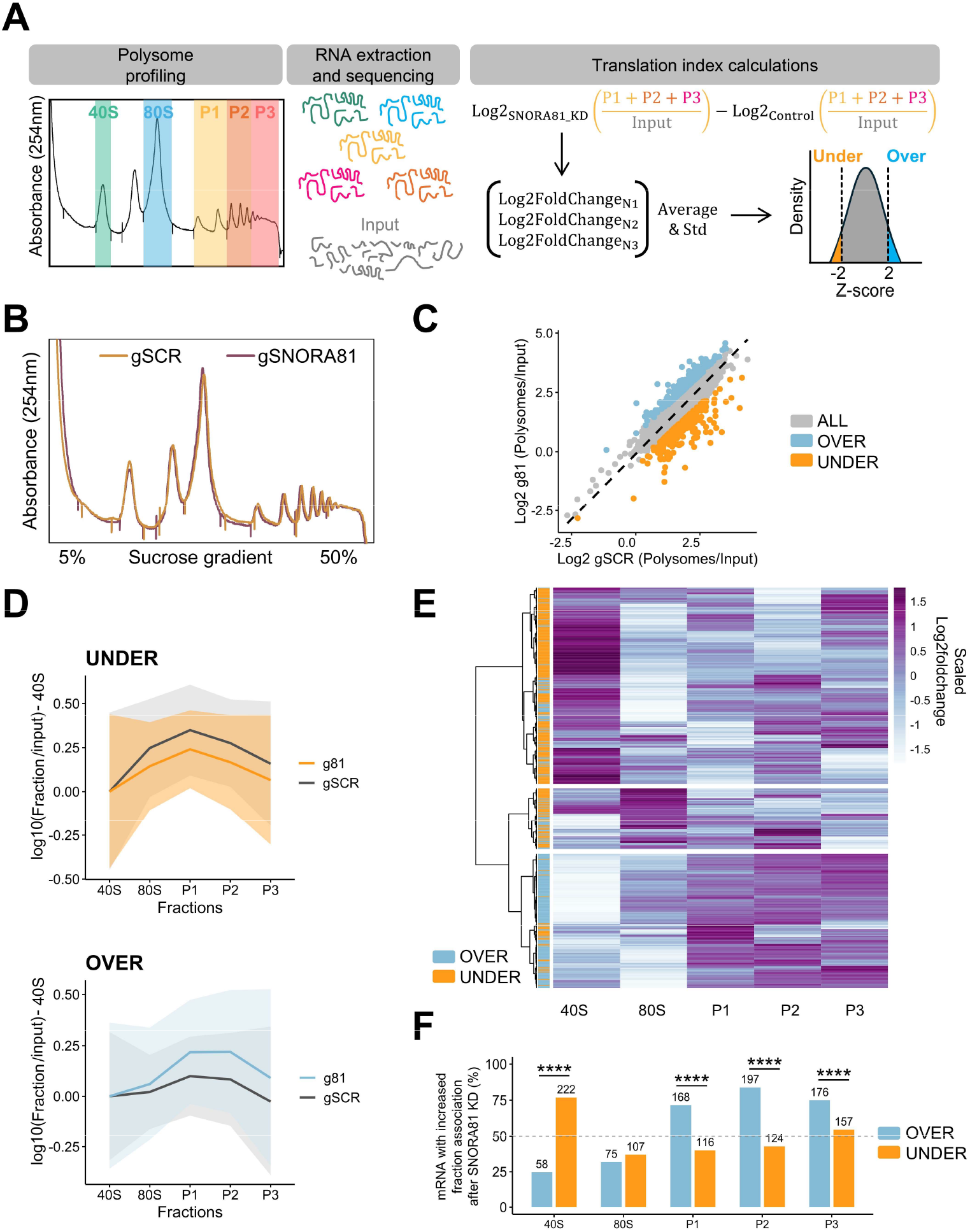
SNORA81-guided Ψ4636 redistributes ribosome association of STAR mRNAs. **(A)** Polysome-profiling workflow. OVCAR-3 cells expressing gSCR (control guide) or gSNORA81 (SNORA81-targeting guides) were subjected to sucrose-gradient fractionation, and RNA from input and gradient fractions was sequenced. Transcripts with ribosome-association z-scores of ≤ −2 or ≥ 2 were classified as under-associated or over-associated SNORA81-dependent translationally altered mRNAs (STAR mRNAs), respectively. Created with BioRender. **(B)** Representative absorbance profiles at 254 nm from gSCR and gSNORA81 OVCAR-3 cells. **(C)** Transcript-level log2 polysome-to-input ratios in gSNORA81 cells versus gSCR controls. Over-associated and under-associated STAR mRNAs are highlighted in blue and orange; non-STAR mRNAs are shown in grey. Dashed line, identity line. **(D)** Mean fraction-distribution profiles of under and over STAR mRNAs across 40S, 80S and polysomal fractions. Values are log10 fraction abundance/input abundance normalized to the 40S fraction. Shaded areas indicate s.d. across n = 3 biologically independent samples. **(E)** Heatmap of scaled log2 fold changes in fraction-specific RNA abundance for individual over-associated and under-associated STAR mRNAs. Rows were hierarchically clustered and row-scaled to highlight redistribution across fractions. **(F)** Fraction-specific redistribution of over and under STAR mRNAs after SNORA81 depletion. Bars show the percentage of STAR mRNAs whose relative representation within each ribosomal fraction increased after SNORA81 depletion (KD/SCR relative representation > 1); numbers indicate transcript counts. For each fraction, the proportion of shifted transcripts was compared within over and under associated mRNAs using chi-square tests (***P < 0.0001; exact P values are provided in methods).

The two STAR groups displayed distinct fraction-specific behaviours (**Figure 3D**). Heatmaps of individual transcripts showed that under-associated STAR mRNAs shifted toward lighter fractions (40S and 80S) after SNORA81 depletion, whereas over-associated STAR mRNAs accumulated in heavier polysomal fractions (**Figure 3E**). Quantification across fractions confirmed distinct redistribution patterns for the under-associated and over-associated STAR groups. For each ribosomal fraction, we calculated the proportion of STAR mRNAs that increased their relative representation following SNORA81 depletion and compared the over-associated and under-associated STAR groups. Under-associated transcripts preferentially shifted toward lighter fractions (40S and 80S), whereas over-associated transcripts preferentially accumulated in polysomes (**Figure 3F**).

To examine ribosome behaviour along these transcripts, we analysed footprint distribution across coding sequences of the 524 STAR mRNAs. Over-associated STAR mRNAs showed relatively increased ribosome occupancy within the first 250 nucleotides of the CDS after SNORA81 depletion, whereas under-associated STAR mRNAs showed the opposite trend (**Figure S8**). These patterns parallel the polysome redistribution in **Figure 3E, F** and indicate transcript-specific changes in ribosome occupancy near the beginning of the CDS. Such changes may reflect altered initiation or early elongation and cannot be distinguished by this analysis alone.

These changes in ribosome association were largely independent of mRNA abundance. Of approximately 13,000 detected protein-coding transcripts, only 292 showed differential abundance upon SNORA81 depletion, with minimal overlap (39 mRNAs) with STAR mRNAs (**Figure S7C**). Changes in mRNA abundance correlated poorly with changes in ribosome association z-score (**Figure S7D**), and both STAR groups displayed relatively low abundance across the transcriptome (**Figure S7E**), further indicating that differential ribosome association is not driven by transcript abundance. Thus, SNORA81 depletion selectively redistributes STAR mRNAs across polysome fractions despite broadly preserved bulk polysome profiles.

### Sequence composition defines STAR mRNAs

We next asked what features distinguish the mRNAs whose ribosome association changes following SNORA81 depletion. Both over- and under-associated STAR mRNAs had significantly shorter coding sequences than all detected mRNAs, and under-associated mRNAs also had shorter 3′ untranslated regions (**Figure S9A**). Both groups contained slightly fewer introns than the overall transcript population (**Figure S9B**). Because these features changed in the same direction in both STAR classes, they do not readily explain their opposite changes in ribosome association following SNORA81 depletion.

We then examined nucleotide composition across transcript regions and found a marked GC bias. Under-associated STAR mRNAs were GC-rich, whereas over-associated STAR mRNAs were AU-rich, particularly within coding sequences and 3′UTRs, with smaller differences in 5′UTRs (**Figure 4A**). Across the transcriptome, CDS GC content showed a modest negative relationship with ribosome association in control cells that significantly strengthened after SNORA81 depletion (Fisher’s r-to-z test, P = 1.16 × 10⁻²⁸) (**Figure 4B**). Thus, SNORA81 depletion strengthens the relationship between coding-sequence composition and ribosome association across the detected transcript population, beyond the STAR mRNA subset. To determine whether this compositional association depended on the threshold used to define STAR mRNAs, we repeated the analysis using alternative z-score thresholds. The GC-content difference between over-associated and under-associated transcripts was maintained across thresholds, indicating that the association between SNORA81-dependent ribosome redistribution and coding-sequence composition is not attributable to a specific STAR threshold (**Figure S9C**).

**Figure 4:**
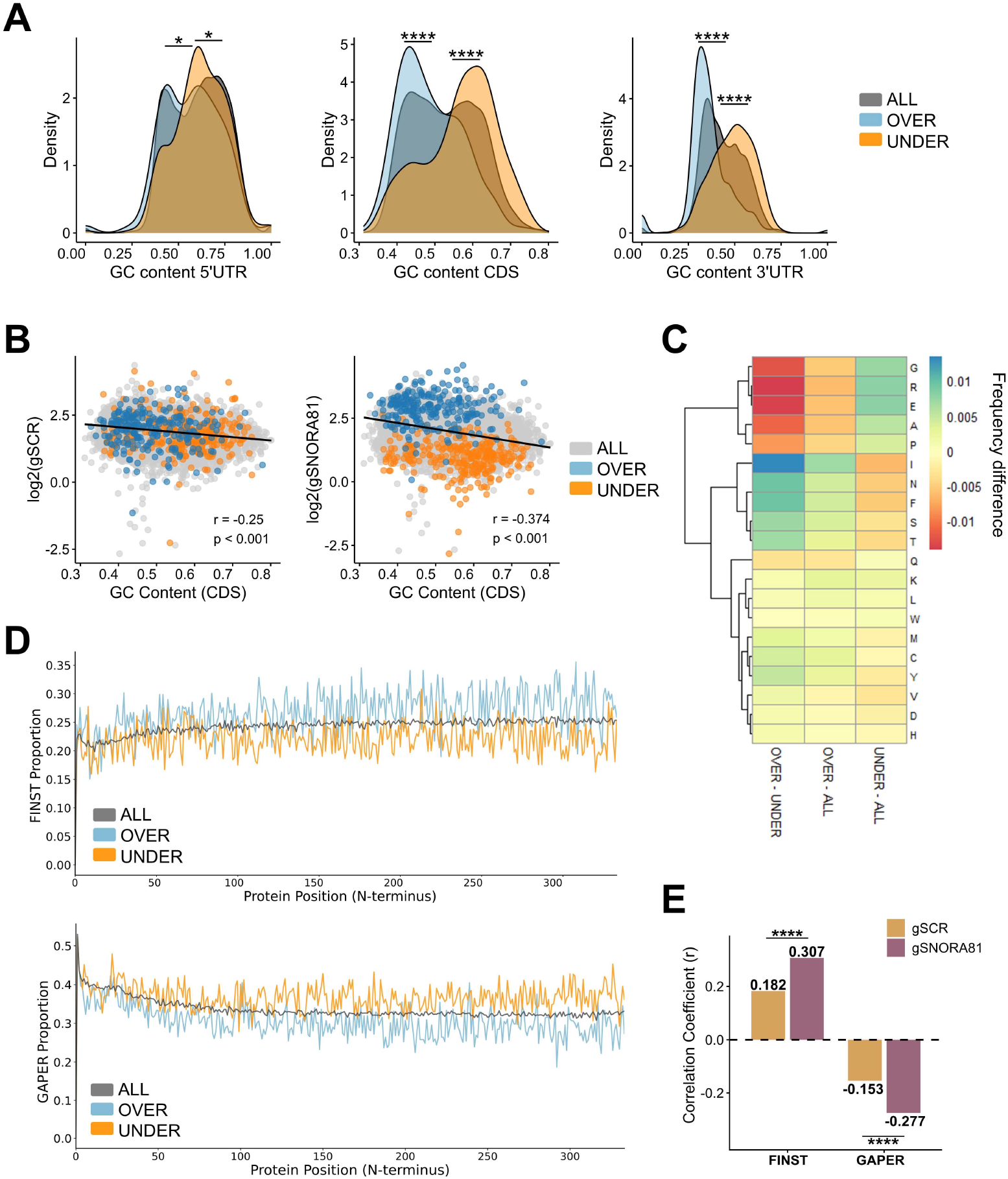
STAR mRNAs differ in GC content and amino-acid composition. **(A)** GC-content distributions in the 5′ UTR, CDS and 3′ UTR for all detected mRNAs (grey; n = 13,025), over-associated STAR mRNAs (blue; n = 235) and under-associated STAR mRNAs (orange; n = 289). STAR mRNAs are SNORA81-dependent translationally altered transcripts with increased or decreased ribosome association after SNORA81 depletion. Significance was assessed by two-sided Kolmogorov–Smirnov tests (*P < 0.05; ****P < 0.0001; exact P values in methods). **(B)** CDS GC content versus ribosome association, defined as polysome enrichment relative to input, in gSCR (control guide) and gSNORA81 (SNORA81-targeting guides) OVCAR-3 cells. Over and under STAR mRNAs are highlighted; black lines show linear fits, with Pearson’s r and P values indicated. Correlations differed significantly (n = 13,025, P = 1.16 × 10⁻^28^, Fisher’s r-to-z test). **(C)** Heatmap of mean amino-acid usage differences for over versus under, over versus all and under versus all mRNAs. Rows are amino acids and hierarchically clustered. Blue indicates higher usage in the first group; red indicates lower usage. **(D)** Positional profiles of FINST and GAPER amino-acid proportions across the first 333 N-terminal residues for all, over and under mRNAs. FINST, phenylalanine, isoleucine, asparagine, serine and threonine; GAPER, glycine, alanine, proline, glutamate and arginine. **(E)** Pearson correlations between transcript-level FINST or GAPER content and ribosome association in gSCR and gSNORA81 cells, across all detected mRNAs (n = 12,984). Dashed line, r = 0. Differences between conditions were assessed by Fisher’s r-to-z test (FINST: P = 7.41 × 10^-27^; GAPER: P = 9.51 × 10^-26^).

Because GC-rich sequences can form stable RNA structures, we examined transcript stability and predicted G-quadruplexes. Reanalysis of published mRNA half-life data^24^ showed that over-associated STAR mRNAs were significantly less stable, whereas the stability of under-associated STAR mRNAs did not differ significantly from that of the overall transcript population (**Figure S9D**). Predicted coding-sequence G-quadruplexes (G4s) were more frequent among under-associated than over-associated STAR mRNAs, with at least one predicted G4 in 55 of 289 and 10 of 235 transcripts, respectively (two-sided Fisher’s exact test, P = 1.14 × 10⁻⁷; **Figure S9E**). Although G4-containing transcripts were enriched among under-associated STAR mRNAs, they represented only a minority of this group and therefore cannot account for most SNORA81-dependent changes in ribosome association. Whether this enrichment is independent of CDS GC content remains unresolved.

Hierarchical clustering of amino-acid frequencies identified two opposing signatures: FINST residues, phenylalanine, isoleucine, asparagine, serine and threonine, occurred at higher frequencies in over-associated STAR mRNAs, whereas GAPER residues, glycine, alanine, proline, glutamate and arginine, occurred at higher frequencies in under-associated STAR mRNAs (**Figure 4C**). Positional analysis across the first 333 N-terminal residues showed that the FINST and GAPER differences were distributed throughout the examined region rather than restricted to discrete positions (**Figure 4D**). Together with the transcript-wide amino-acid frequencies, this positional analysis associates SNORA81-sensitive ribosome behaviour with broad coding-sequence composition rather than a single local motif. These observations prompted us to ask whether the amino-acid biases were linked to ribosome association. In control cells, FINST frequency correlated positively with ribosome association, whereas GAPER frequency correlated negatively; both correlations changed significantly after SNORA81 depletion (FINST, P = 7.41 × 10⁻²⁷; GAPER, P = 9.51 × 10⁻²⁶; Fisher’s r-to-z tests; **Figure 4E**).

To determine whether these compositional features were independently associated with STAR classification, we performed multivariate logistic regression including CDS GC content, FINST and GAPER frequencies, transcript-region lengths, mRNA abundance and codon adaptation index. Over-associated STAR mRNAs were independently characterized by lower CDS GC content and higher FINST amino-acid frequency, whereas under-associated STAR mRNAs were independently associated with higher CDS GC content and lower FINST frequency (**Figure S10A**). Together, these findings define a transcript-composition signature associated with SNORA81-dependent changes in ribosome association, comprising CDS GC content and codon and amino-acid features.

### STAR mRNAs converge on cancer-associated programs

Having identified a transcript-composition signature associated with STAR mRNAs, we asked whether this selective redistribution is reflected in cancer-associated protein outputs. Gene Ontology enrichment analysis of under-associated and over-associated STAR mRNAs identified cancer-relevant processes, including signalling and gene regulation, developmental and differentiation programs, proliferation and cell-cycle control, migration and invasion, and tumour microenvironment-associated pathways (**Figure 5A**). CancerMine annotation identified cancer-associated genes within both STAR-mRNA groups (**Figure 5A**). These enriched processes overlapped with pathways identified in complementary analyses of SNORA81 depletion, including PI3K–AKT signalling, inflammatory and stress-response pathways, and regulators of proliferation and migration^25^. This overlap supports the functional coherence of the cancer-associated mRNAs whose ribosome association changes following SNORA81 depletion.

**Figure 5:**
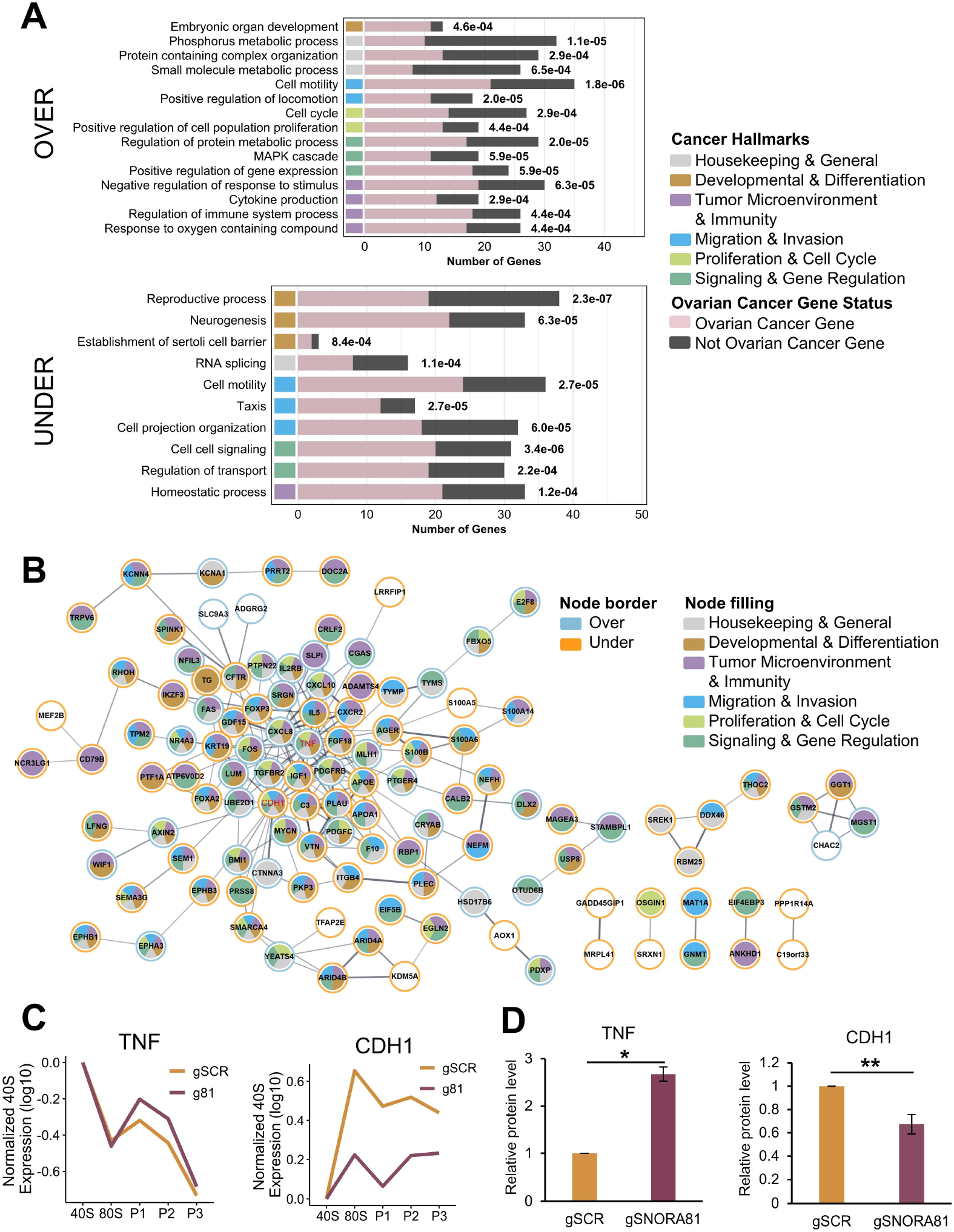
SNORA81-guided Ψ4636 alters translation of cancer-associated mRNAs. **(A)** Gene Ontology biological-process enrichment of over and under-associated SNORA81-dependent translationally altered mRNAs (STAR mRNAs). Redundant terms were collapsed; top non-redundant terms with FDR < 0.001 are shown. Terms were assigned to cancer-hallmark-related categories: housekeeping and general, development and differentiation, tumour microenvironment and immunity, proliferation and cell cycle, migration and invasion, and signalling and gene regulation. Bars indicate STAR mRNAs counts; labels indicate gene ratio and FDR. Cancer-associated genes, defined using CancerMine, are highlighted in pink. **(B)** STRING protein-protein interaction network of CancerMine-associated STAR mRNAs. The network includes 118 proteins with at least one interaction. Nodes are manually colored by cancer hallmark-related gene ontology category from a; borders indicate under-associated (orange) or over-associated (blue) STAR mRNAs. The network shows significant protein-protein interaction enrichment (P = 6.61× 10^-14^). **(C)** Validation of selected STAR mRNAs at the level of ribosome association after SNORA81 depletion. Line plots show input-normalized, 40S-corrected fraction distributions for TNF (over-associated), and CDH1 (under-associated), across 40S, 80S and polysomal fractions. Values are log10 fraction abundance/input abundance for gSCR (control guide) and gSNORA81 (SNORA81-targeting guides) cells. **(D)** Validation of translational changes at the protein level after SNORA81 depletion. Bar plots show relative TNF and CDH1 protein abundance measured by immunoblotting 48 hours (TNF) or 72 hours (CDH1) after transfection with gSCR or gSNORA81. Protein levels were normalized to GAPDH and scaled to gSCR = 1. Data are mean ± s.d. from n = 3 biologically independent experiments; P values were determined by two-sided t-tests, (P = 0.004 **, P = 0.04 *).

To determine whether the proteins encoded by cancer-associated STAR mRNAs form a connected network, we performed STRING protein–protein interaction analysis of CancerMine-associated STAR mRNAs. Of these proteins, 118 had at least one STRING interaction and formed a significantly interconnected network (PPI enrichment P = 6.61 × 10⁻¹⁴; **Figure 5B**), with interactions spanning multiple cancer-hallmark-related biological processes, including proliferation, signaling, migration and invasion, and tumour microenvironment-associated pathways. These results show that proteins encoded by CancerMine-associated STAR mRNAs form a functionally connected network rather than a collection of isolated gene products.

We next asked whether pathway-level changes in STAR mRNAs were reflected at the protein level. To prioritize candidates for validation, we selected tumour necrosis factor (TNF) and E-cadherin (CDH1) because they were among the most highly connected over-associated and under-associated STAR mRNAs, respectively, within the STRING network (red nodes in **Figure 5B**). In the same direction as its increased polysome association (**Figure 5C**), TNF protein abundance increased after SNORA81 depletion (**Figure 5D; Figure S10B**). Because TNF is a central inflammatory and stress-response regulator implicated in ovarian cancer biology^26,27^, this result provides one protein-level example consistent with the enrichment of inflammatory, stress-response and tumour microenvironment-associated functions among STAR mRNAs. Conversely, decreased polysome association of CDH1 mRNA (**Figure 5C**) was accompanied by reduced E-cadherin protein abundance after SNORA81 depletion (**Figure 5D; Figure S10B**). We interpret CDH1 primarily as an example in which altered ribosome association is accompanied by a corresponding change in protein abundance, rather than as a standalone marker of invasive behaviour, because E-cadherin has context-dependent roles in cancer progression, including both loss during invasion and re-expression in metastatic or proliferative states^28,29^. Although loss of E-cadherin is commonly associated with epithelial–mesenchymal transition and invasion, E-cadherin can be retained or re-expressed in some ovarian tumours and metastases^30,31^. As an additional example, we examined RBM25, another under-associated STAR mRNA, and observed reduced RBM25 protein abundance following SNORA81 depletion (**Figure S10B**). Collectively, the pathway enrichment and representative protein changes support a model in which increased SNORA81 abundance in HGSC increases pseudouridylation at U4636, thereby increasing the fraction of ribosomes carrying Ψ4636 within the cellular ribosome pool (**Figure 6**). Together, these data support a model in which SNORA81-dependent pseudouridylation at U4636 may influence the local eL31-proximal ribosomal interface, contributing to changes in ribosome pausing and STAR-mRNA polysome association, with corresponding protein-abundance changes in selected examples.

**Figure 6:**
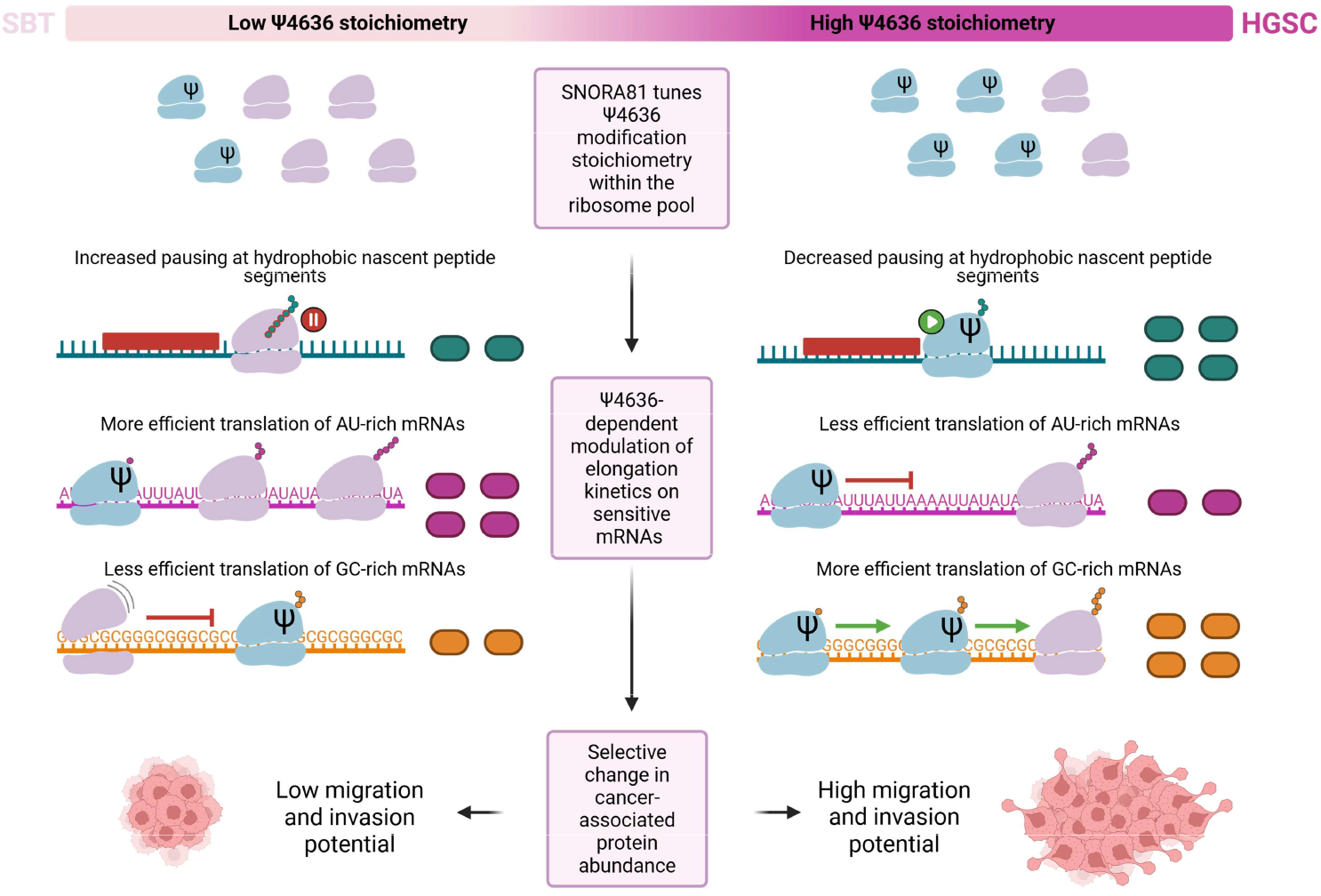
SNORA81-guided Ψ4636 drives selective translation in ovarian cancer through an eL31-proximal ribosomal interface. Proposed model for SNORA81-dependent translational control in ovarian cancer. Elevated SNORA81 increases pseudouridylation at U4636 within the cellular ribosome pool. Ψ4636 occupies a local rRNA–eL31 interface near the peptide-exit region and is proposed to influence codon- and nascent-peptide-associated pausing. These elongation changes redistribute STAR mRNAs across polysomes and alter cancer-associated protein outputs, supporting a model in which modulation of a single snoRNA-guided rRNA modification contributes to translational programs associated with ovarian cancer proliferation, migration and invasion. Ribosomes depicted on mRNAs are shown schematically to illustrate the proposed model and do not represent experimentally determined ribosome positions or densities. Created with BioRender.

## DISCUSSION

Ribosomes were long viewed as invariant machines, yet variation in ribosomal protein content, rRNA sequence and rRNA modifications can bias translation toward distinct mRNA subsets^2–6,32^. Here we identify a mechanism by which a single variable rRNA pseudouridine couples ribosome heterogeneity to cancer-associated translation. SNORA81 depletion reduces pseudouridylation at U4636, a partially modified 28S rRNA site positioned near eL31, and alters ribosome occupancy associated with codon identity and nascent-peptide properties. The position of Ψ4636 at the rRNA-eL31 interface near the peptide-exit region provides a plausible structural basis for this effect.

These changes in ribosome occupancy accompany transcript-level differences in ribosome association, including redistribution of STAR mRNAs across polysomes and corresponding changes in the abundance of representative cancer-associated proteins. Thus, rRNA-modification stoichiometry can act as a regulatory variable linking a local ribosomal interface to selective translation in high-grade serous ovarian cancer.

Foundational studies of rRNA pseudouridylation showed that broad disruption of dyskerin, the pseudouridine synthase responsible for H/ACA snoRNA-guided rRNA modification, perturbs ribosome function, including translational fidelity and IRES-dependent initiation^33,34^. More recent work has established that individual snoRNA-guided rRNA modifications can also shape selective translation. Regulation of site-specific 2’-O-methylation generates heterogeneous ribosome populations that selectively influence translation of distinct mRNA classes^12^, while loss of SNORA24-guided 18S pseudouridine sites alters decoding fidelity, miscoding and stop-codon readthrough^11^. Our findings build on this framework by showing that variation in pseudouridylation at a single 28S rRNA site near the peptide-exit region can influence translational selectivity, with changes in ribosome occupancy that point to altered elongation dynamics.

Mechanistically, U4636 differs from decoding-center sites such as those guided by SNORA24: it lies in H96 at an rRNA–eL31 interface near the peptide-exit region and within a structural framework extending through the H94–H97 junction towards the exit tunnel. Because pseudouridine can stabilize local RNA geometry, and because nascent-peptide properties can regulate elongation and pausing within the exit tunnel, we propose that Ψ4636 influences peptide-dependent elongation indirectly rather than through direct contact with the nascent chain. By influencing the local eL31-proximal interface, Ψ4636 may alter ribosome sensitivity to nascent-peptide properties during elongation^35–37^. The position of H96 relative to eL31, uL3 and the uL4/uL22 constriction provides a plausible structural route through which modification at U4636 could influence elongation. Consistent with the potential functional importance of such local ribosomal interfaces, methylation of uL3 has been shown to alter its interaction with 28S rRNA and modulate codon-dependent translation elongation, providing precedent for modification-dependent changes in ribosomal protein–rRNA interactions to influence elongation^38^. The eL31 interface is notable because eL31 itself has been linked to proliferation and migration in colorectal and gastric cancers^39,40^, but whether modification-dependent interactions at U4636 contribute to these cancer-related functions of eL31 remains unknown^41,42^.

The mRNAs most sensitive to reduced U4636 pseudouridylation segregate into two oppositely regulated classes with distinct GC content, codon usage and amino-acid composition, consistent with a ribosome-filter model in which ribosome state biases translation toward transcripts sharing global features rather than discrete motifs^43,44^. This bidirectional behaviour is important because it argues against a nonspecific loss of translational capacity and instead points to a selective redistribution of ribosome engagement across the transcriptome. Changes in ribosome association are largely independent of changes in transcript abundance, and the affected mRNAs encode proteins involved in proliferation, migration and stress-response pathways. Notably, these cancer-associated pathways overlap with those identified and validated following SNORA81 depletion, including PI3K-AKT signaling and regulators of proliferation and migration^25^. Together, these findings link variation at a single rRNA modification site to selective translational remodeling of cancer-associated programs.

Taken together, our findings recast partial rRNA modification as a regulatory variable rather than a fixed structural feature. Variation in the stoichiometry of a single pseudouridine can bias ribosome pausing and association towards defined mRNA groups without broadly disrupting ribosome biogenesis or bulk polysome organization. We propose that mRNAs with different coding-sequence compositions and nascent-peptide properties have different elongation kinetics on Ψ4636-containing and U4636-containing ribosomes. In this model, mRNA-specific differences in elongation are amplified into changes in ribosome occupancy and polysome association, redistributing transcripts whose coding sequences share GC and amino-acid biases. We term this proposed multilevel cascade the “sno-balling effect”: a change in the abundance of a single snoRNA propagates through rRNA-modification stoichiometry, ribosome function, selective mRNA translation and protein output to reshape cancer-associated pathways and cellular behaviour. In high-grade serous ovarian cancer, elevated SNORA81 and increased pseudouridylation at U4636 therefore provide a route by which malignant cells may establish a selective translational state that reinforces cancer-associated behaviour.

More broadly, our findings identify individual rRNA modifications as potential regulatory nodes linking ribosome heterogeneity to selective translation. Our findings suggest that variation in rRNA-modification stoichiometry can provide a tunable layer of translational control, whereby local changes in ribosome architecture influence elongation and ultimately bias ribosome engagement across mRNA populations. This framework raises the possibility that snoRNA-guided rRNA modification could represent a previously underappreciated layer of translational regulation in cancer and could be explored in future therapeutic studies.

## Limitations of the study

Mechanistic experiments were performed in OVCAR-3 cells. Elevated SNORA81 abundance and U4636 pseudouridylation in primary HGSC samples support the disease relevance of this axis, but the tumour comparison was descriptive and based on a small cohort, so generality across ovarian cancer subtypes awaits validation in additional models and patient-derived systems. SNORA81 depletion reduces rather than eliminates U4636 pseudouridylation, so our experiments report the behavior of mixed ribosome populations; the guide-region rescue supports a modification-dependent contribution to the growth phenotype but does not exclude additional consequences of SNORA81 depletion. Structural analysis establishes contacts between Ψ4636 and eL31, and ribosome-occupancy changes track codon identity and nascent-peptide properties; the functional contribution of this interface and the underlying elongation kinetics remain to be tested directly. Polysome association reports ribosome engagement rather than synthesis rate, and protein-level changes were confirmed for selected targets. Finally, time points were chosen to separate early translational changes from later phenotypic consequences, but some later effects may include secondary responses to SNORA81 depletion. These caveats bound the generality of our conclusions rather than the central observation that a single snoRNA-guided rRNA modification selectively shapes ribosome behavior and cancer-associated translation.

## RESOURCE AVAILABILITY

## Lead contact

Requests for further information and resources should be directed to Sherif Abou Elela.

## Materials availability

Plasmids generated in this study are available from the lead contact upon request.

## Data and Code Availability

• Sequencing data generated in this study have been deposited in the NCBI Gene Expression Omnibus (GEO; https://www.ncbi.nlm.nih.gov/geo) under the accession number for the polysome sequencing GSE341824 and for the footprinting sequencing GSE343641.

• Nanopore raw data were deposited on Zenodo.

• Any additional information required to reanalyze the data reported in this paper is available from the lead contact upon request.

## ACKNOWLEDGEMENTS

The authors are grateful to Homa Ghalei and Sébastien Durand for their thorough review of the manuscript and insightful suggestions. We are also thankful to Eric Westhof, who initially modelled the interaction between eL31 and Ψ4636 and shared valuable insights and discussion regarding the local environment of this modification. We also thank the members of the Abou Elela and Scott laboratories for valuable discussions. Finally, we acknowledge the Digital Research Alliance of Canada for providing the high-performance computing infrastructure that supported this work.

This work was supported by the Canadian Institutes of Health Research (CIHR) grant 486207 (S.A. and M.S.S.). S.A. holds a Canada Research Chair in RNA Biology and Cancer Genomics. M.S.S. holds a Canada Research Chair in Bioinformatics of Non-Coding RNA. LFG was awarded scholarships from the FRQS and from the NSERC CREATE fund as part of the RNA innovation program. B.P.K. acknowledges support from Institut National du Cancer (INCa), the Fondation pour la Recherche Médicale (FRM) and EquipEx+ France-Cryo-EM (ANR-21-ESRE-0046).

## AUTHOR CONTRIBUTIONS

L.F.-G. designed and performed experiments. A.R. performed western blot experiments. B.S.d.P., É.F.-C. and G.D.-F. analysed sequencing data. V.M and Y.M carried out the HydraPsiSeq experiment. C. B., B. P.K. modelled the interaction between eL31 and Ψ4636. T.A. helped establish the CRISPR–Cas13d system. M.S.S. designed study concepts, supervised data analysis, and reviewed and edited the manuscript. S.A.E. conceived and supervised the study. L.F.-G. and S.A.E. wrote the manuscript with input from all authors.

## SUPPLEMENTARY INFORMATION

Document S1. Supplemental Figures S1-S10 and Tables S1-S2

## STAR★METHODS

### Key resources table

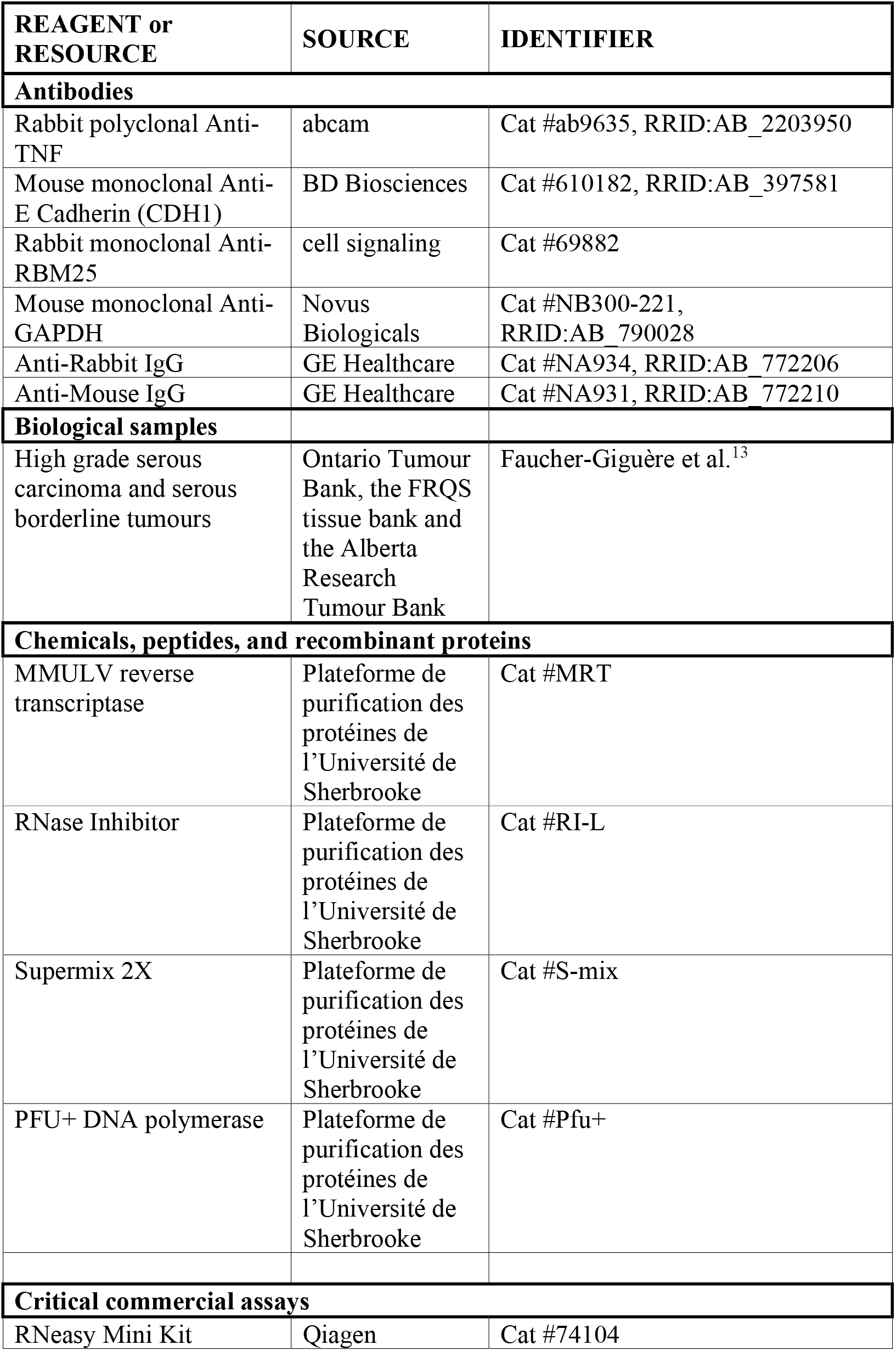
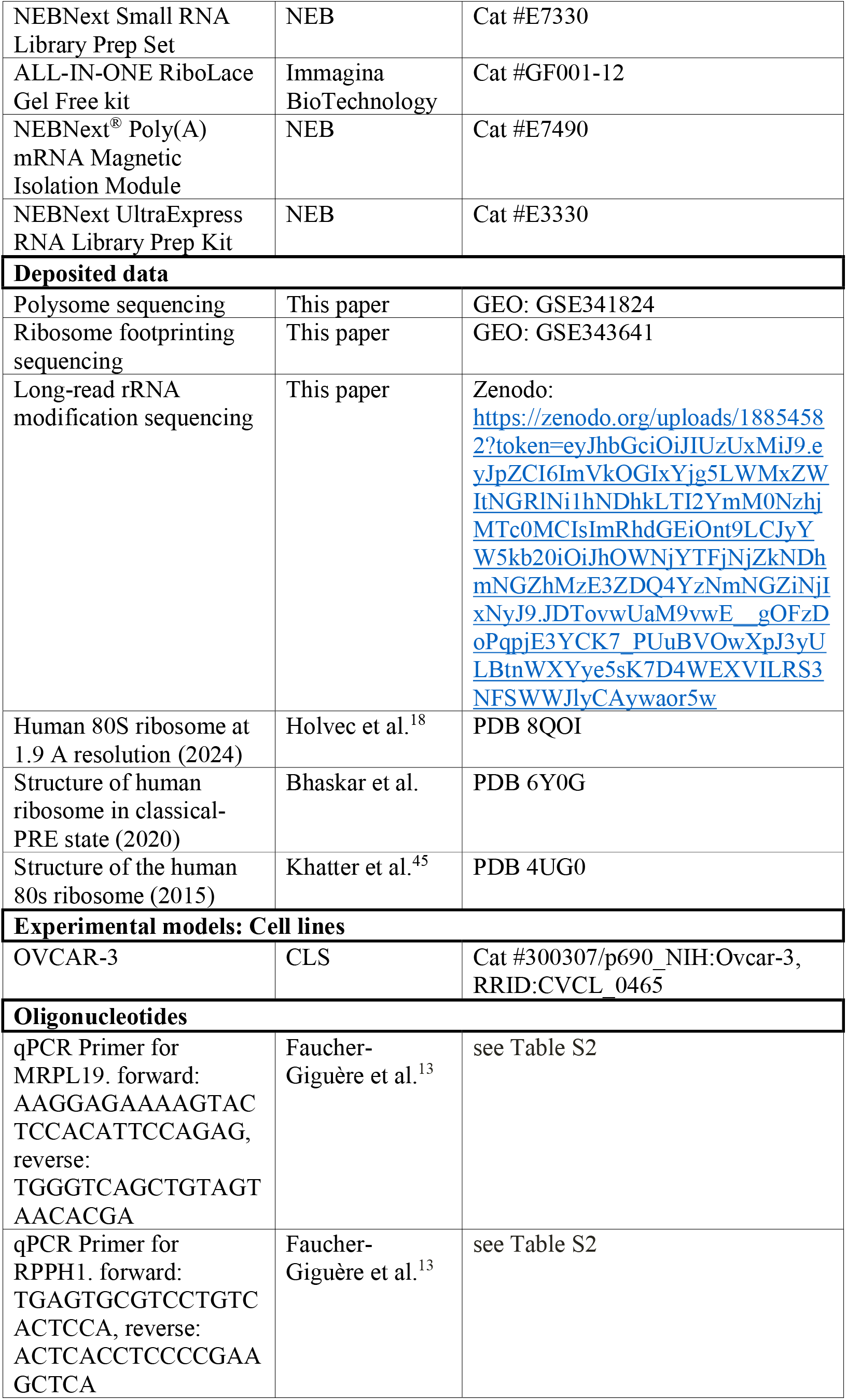
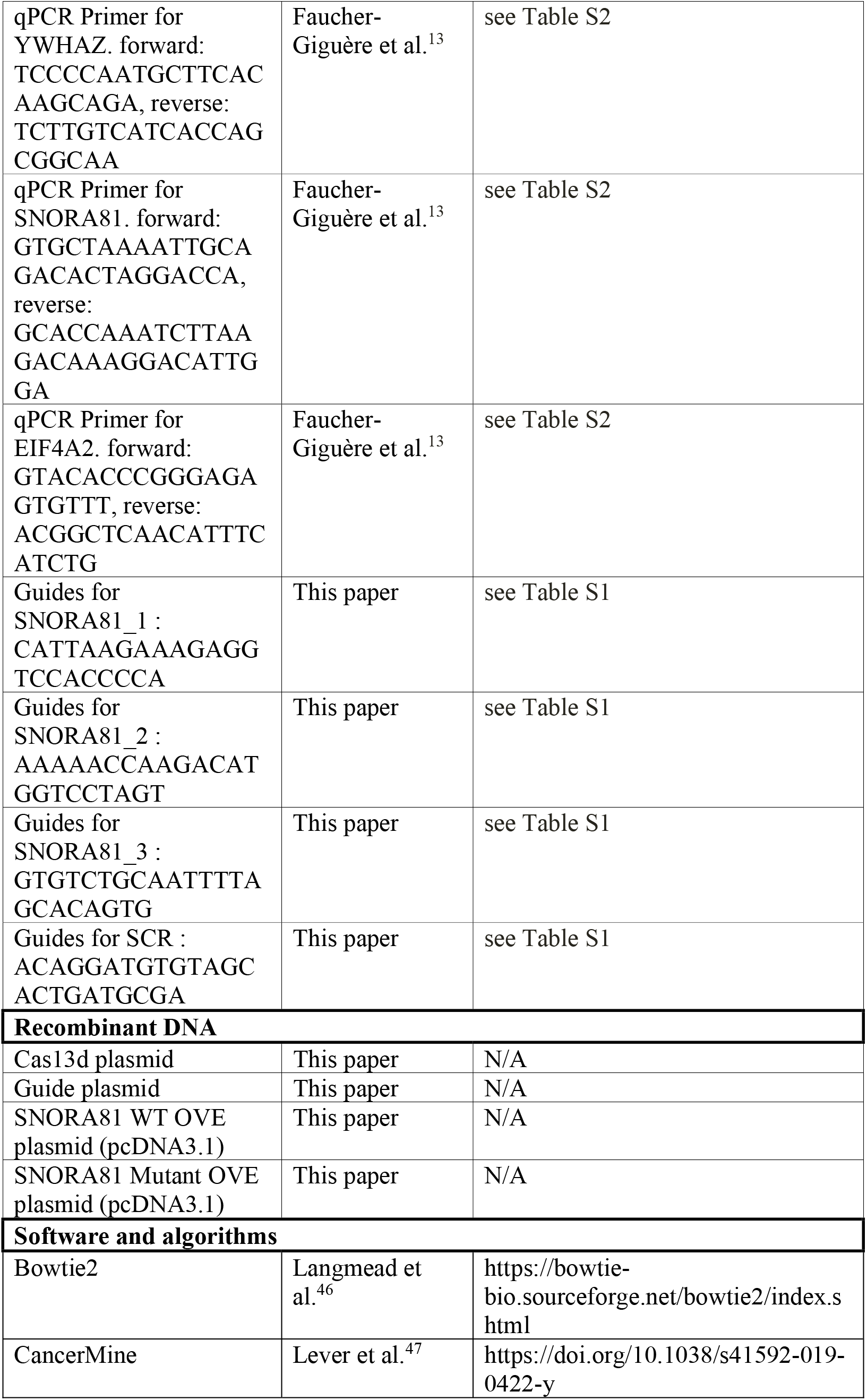
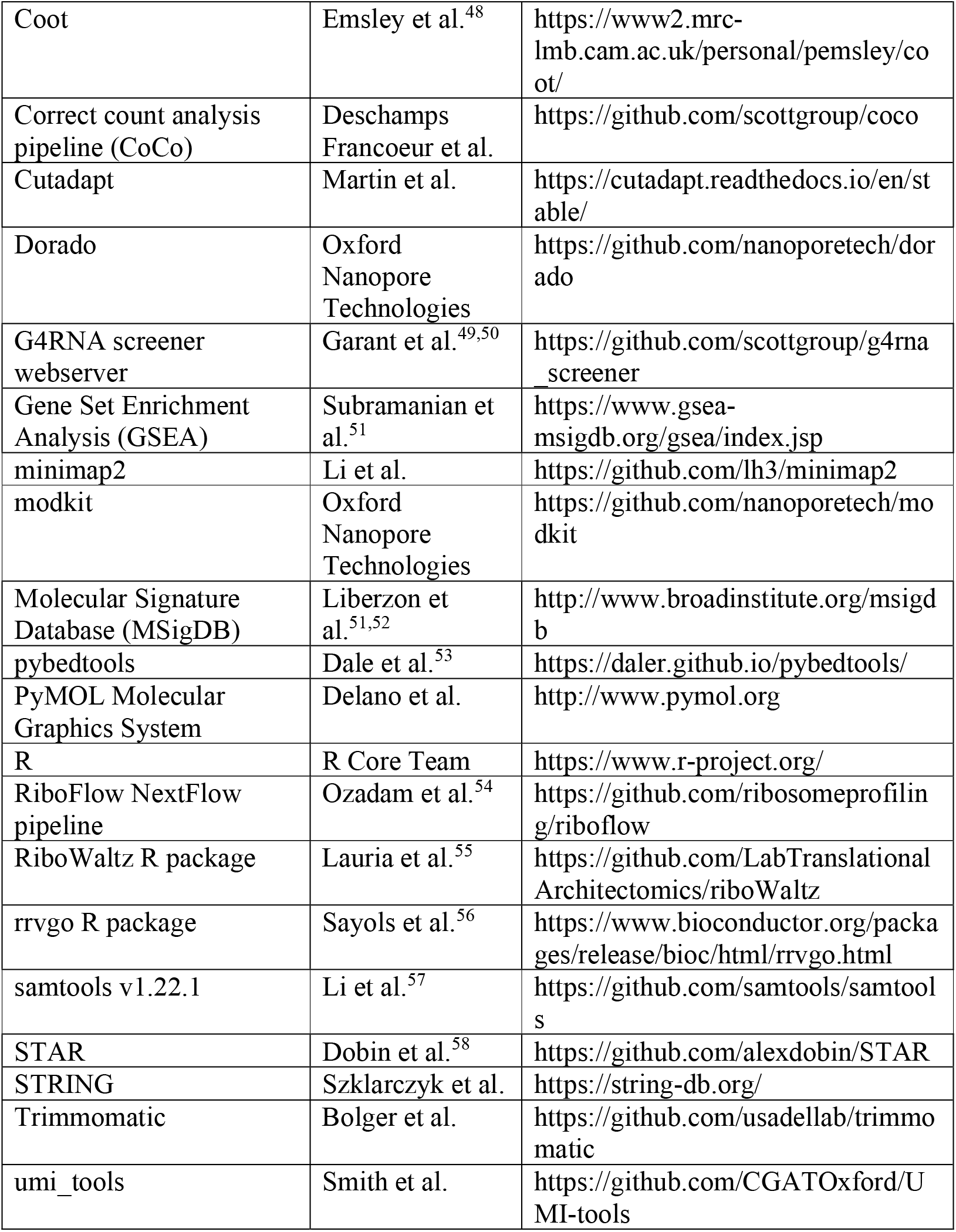

### Experimental model and study participant details

#### Human tissue samples

Sequencing of High-grade serous ovarian carcinoma (HGSC) and Serous borderline tumours (SBT) from female sex, no report of gender. HGSC and SBT tissues were provided by the Ontario Tumour Bank, the FRQS tissue bank and the Alberta Research Tumour Bank. Patient information and consent were managed by the tumour banks and released for this study under their respective material transfer agreements. Tissue handling and experimental procedures were approved by the Comité d’éthique de la recherche du CIUSSS de l’Estrie–CHUS (CÉR #10-062, 2010-241). More information about the provenance and quality control of the tissues was previously described^13^.

#### Cell lines

OVCAR-3 ovarian cancer cells (CLS, cat. no. 300307/p690_NIH:OVCAR-3) were cultured in RPMI 1640 medium supplemented with 20% fetal bovine serum (FBS) and 2 mM L-glutamine. Cells were maintained at 37°C in a humidified incubator with 5% CO₂ and passaged according to ATCC guidelines. Cell lines were used for no more than 20 passages. Mycoplasma testing was performed routinely by qPCR through the Université de Sherbrooke RNomics platform.

### Method details

#### Cell culture and CRISPR-Cas13d-mediated SNORA81 depletion

For CRISPR–Cas13d-mediated SNORA81 depletion, cells were transfected with a Cas13d expression plasmid and guide RNA plasmids using Lipofectamine 2000 (Invitrogen) according to the manufacturer’s instructions. Cells were seeded 24 hours before transfection at 3.0 × 10⁵ cells per well in six-well plates or 1.0 × 10⁶ cells per 10 cm dish. For six-well plates, cells were transfected with 1 µg total plasmid DNA, comprising 500 ng Cas13d plasmid and 500 ng guide plasmid. For 10 cm dishes, cells were transfected with 4 µg total plasmid DNA, comprising 2 µg Cas13d plasmid and 2 µg guide plasmid. Guide plasmids encoded either a scrambled control guide or a pool of three guides targeting SNORA81. Guide RNA sequences are listed in **Table S1.**

#### RNA extraction, cDNA synthesis and RT–qPCR

Total RNA was extracted as described previously^13^, using the RNeasy Mini Kit (Qiagen) with on-column DNase digestion (RNase-Free DNase Set, Qiagen). To improve recovery of small RNAs, 1.5 volumes of 100% ethanol were added during the precipitation step. RNA concentration and integrity were assessed using a 2100 Bioanalyzer (Agilent). For cDNA synthesis, 500 ng of total RNA was reverse transcribed using random hexamers and MMULV reverse transcriptase. Enzyme for reverse transcription (#MRT, #RI-L) and Supermix 2X (#S-mix) for qPCR were purchased from the Plateforme de purification des protéines de l’Université de Sherbrooke. Relative transcript levels were quantified by RT-qPCR and calculated using the comparative ΔΔCq method after normalization to a panel of reference genes. No-template and no-reverse-transcriptase controls were included as negative controls in all experiments. Statistical significance between conditions was assessed using a two-sided Student’s t-test with unequal variance. Primer sequences are listed in **Table S2**.

#### Xcelligence assay

xCELLigence assays were performed as described previously^13^. Briefly, 50 µl complete medium was added to each well of an E-Plate and used for background measurement. OVCAR-3 cells were then seeded at 5,000 cells per well and monitored until the Cell Index reached approximately 0.6 (typically 24–48 hours after seeding). Cells were then transfected with 50 µl transfection mixture containing the indicated Cas13d and guide RNA plasmids. Plates were maintained at 37°C with 5% CO₂, and impedance-based Cell Index measurements were recorded every 10 min.

For the rescue experiments, OVCAR-3 cells were first transfected with the CRISPR-Cas13d together with either SNORA81-targeting or control (gSCR) guide RNAs. 24 hours later, cells were transfected with plasmids expressing either wild-type (WT) SNORA81 or a guide-region mutant SNORA81. Rescue plasmids were generated by PCR amplification of the EIF4A2 genomic region spanning exons 7-8, including the intron containing SNORA81, followed by cloning into the pcDNA3.1 vector. The mutant construct contained a shuffle SNORA81 guide sequence but preserving the overall snoRNA structure.

#### CLAP analysis of U4636 pseudouridylation

Single site pseudouridylation at U4636 was quantified by CLAP (CMC-RT and ligation assisted PCR analysis of pseudouridylation modification) as previously described^13^. Total RNA was collected 48 hours after transfection with Cas13d and either SNORA81-targeting or scrambled control guide plasmids. Briefly, total RNA was treated with CMC prior to reverse transcription, adaptor ligation and PCR amplification of the target region. Modified and unmodified products were resolved and quantified to determine the fraction of pseudouridylated transcripts. Unless otherwise indicated, the protocol was performed like previously described^13^. PFU7 DNA polymerase produced by the Université de Sherbrooke Protein Purification platform was used for PCR amplification in place of the commercial enzyme used in the original protocol^15^.

#### HydraPsiSeq

Global pseudouridylation levels were quantified by HydraPsiSeq as previously described^14^. Briefly, 300 ng of total RNA was subjected to hydrazine/aniline cleavage, converted into sequencing libraries using the NEBNext Small RNA Library Prep Set for Illumina (NEB E7330), and sequenced on an Illumina NextSeq 2000 platform (50-bp single-end reads). Sequencing reads were processed using the published HydraPsiSeq pipeline^14^, including adapter trimming, alignment to human rRNA sequences, and calculation of HydraPsiSeq scores from normalized cleavage profiles.

#### Direct RNA modification analysis

Long-read nanopore sequencing was performed 48 hours after SNORA81 depletion to allow turnover of pre-existing modified rRNA and detection of steady-state changes in pseudouridylation. 500 ng RNA was polyadenylated like previously described^59^. RNA was barcoded and sequenced by the Université de Sherbrooke RNomics platform.

Analysis of long-read data was performed as previously described^59^. Briefly, raw POD5 files were basecalled using Dorado (v1.3.0) and aligned to a pool of ribosomal RNA variants with minimap2 (v2.3). Pseudouridylation frequency at each residue was predicted using the Dorado model rna004_130bps_sup@5.1.0 on samples with a minimum read depth of 20,000. Sorting and indexation of the bam files was done using samtools v1.22.1^57^. The files were then fed to modkit pileup v0.5.1 https://github.com/nanoporetech/modkit to extract and to compile pseudouridylation events across each uridine base with the options “--motif T 0 --filter-threshold T:0.90”. Only pseudouridine sites validated in the union of two datasets^18^ were retained. Additionally, predictions were filtered to include only positions with a minimum local coverage of 20 reads and a modification frequency of at least 15%.

#### Structural analysis

Structural analyses were performed using Coot^48^ and PyMOL software (http://www.pymol.org). The human 80S ribosome structure PDB 8QOI was used for the main structural representations of the Ψ4636-containing ribosome^18^. Hydrogen bonds between nucleotide Ψ4636 and eL31 residues were identified by structural inspection in Coot, considering distances and geometry. To position conserved functional centres, ribosome structures were aligned with PDB 6Y0G, which was used to map the mRNA and tRNA positions. The peptide exit tunnel was visualized by mapping the tunnel path and coordinates reported for the human ribosome structure PDB 4UG0 ^45^ in a previous structural analysis^60^.

#### RNA integrity and rRNA analysis

Mature 28S and 18S rRNA levels and the 28S/18S ratio were quantified across multiple time points after SNORA81 depletion, consistent with the multi-day half-life of mature ribosomes. Pre-rRNA processing intermediates were assessed by northern blotting using probes against 5′ETS, ITS1, ITS2 and 3′ETS regions^61,62^ processing and main steps were done as previously described^63^.

#### Ribosome Profiling and P-site assignment

Ribosome profiling was performed using the ALL-IN-ONE RiboLace Gel Free kit (Immagina BioTechnology) according to the manufacturer’s instructions. Briefly, actively translating ribosomes were enriched from OVCAR-3 cells 24 hours after SNORA81 depletion. Matched polyA-selected input RNA libraries were prepared as described below (see RNA-sequencing section); whereas ribosome-protected fragments (RPF) were prepared according to the manufacturer’s protocol. RPF libraries were sequenced on an Element AVITI System using a Cloudbreak Freestyle High Output 150 cycles kit (820-00022) (1X154). Sequencing reads were first aligned to a contaminant reference containing rRNA, tRNA and other non-coding RNA sequences using Bowtie2^46^ and matching reads were discarded. The remaining reads were aligned to the human genome (hg38, v101) using STAR^58^ and default parameters. Reads mapping to protein-coding genes, long non-coding RNAs and pseudogenes were retained for downstream analysis. Transcriptome-aligned BAM files were generated using the --quantMode TranscriptomeSAM option and used for downstream ribosome profiling analyses.

Ribosome-profiling data were analysed using the RiboWaltz R package (2.0)^55^ to determine P-site offsets and quantify codon-resolved ribosome occupancy. Only RPFs of 28-34nt were retained for downstream analysis (length_filter_mode = “custom” and length_range = 28:34). P-site offsets were estimated using default parameters (flanking = 6) with automatic selection of read extremity (extremity = “auto”), which consistently selected the 5’end across all retained read lengths. Quality control included assessment of read-length distributions, coding-sequence enrichment, transcript coverage, reading-frame periodicity and metagene profiles around annotated start and stop codons. QC summaries are shown in **Figure S5**.

#### Ribosome pausing analysis

To characterize ribosome pausing, ribosome footprinting reads were processed using the RiboFlow NextFlow pipeline^54^. Briefly, unique molecular identifiers (UMIs) were extracted using umi_tools with a 4nt dual UMI structure (5’ and 3’ end). Reads were adapted-trimmed using Cutadapt (adapter sequence: TCTCCTTGCATAATCACCAACC; minimum overlap 4nt; quality-cutoff: Phred 28). Reads were aligned to the human transcriptome (APPRIS human v2 principal isoform). Processed outputs were stored as .ribo files for downstream analysis.

Pause sites were identified as previously described^64^. Briefly, per-nucleotide ribosome occupancy within coding sequences was modelled using a negative binomial distribution, and significance was assessed across biological replicates. Replicate p-values were combined using Fisher’s method, and sites with a combined P ≤ 10⁻⁶ were retained.

#### Physicochemical properties of ribosome exit tunnel regions

Condition specific pause sites were identified from the full pause-site dataset (gSNORA81-specific n=1354; gSCR-specific n=1001). For physiochemical analyses pause sites were retained only if they could be mapped to protein-coding Ensembl id and corresponding protein sequences, yielding 1192 KD-specific and 898 gSCR-specific sites with extractable 30 amino acid upstream the tunnel sequences. For each pause site, the upstream amino-acid sequence spanning the ribosome exit tunnel (-30 to -1 relative to p-site) was extracted from the human reference genome (hg38, APPRIS principal isoforms). Amino-acid residues were mapped to physiochemical properties including charge, hydrophobicity (Kyte–Doolittle scale), polarity and volume. Position-resolved property scores were computed separately for SNORA81 knockdown (gSNORA81) and control (gSCR) conditions. For each position within the tunnel window, property distributions between conditions were compared using two-sided Mann-Whitney U tests. Mean property values and standard error of the mean (SEM) were computed across all pause sites.

#### Tunnel radius calculation

Tunnel geometry was extracted from the 8QOI ribosome structure using previously mapped tunnel centerline coordinates from 4UG0 model. Tunnel radius was computed in PyMOL as the minimum Euclidean distance between each tunnel centerline point and the ribosomal atomic structure, corresponding to the shortest distance from the tunnel centerline to surrounding ribosomal atoms. This approach is consistent with previously described geometric analyses of ribosome exit tunnels^60^. The tunnel coordinates (x, y, z), index ordering, and radius values were then exported for downstream analysis performed in R. The radius profile was smoothed (R smooth.spline, spar = 0.5) to reduce local fluctuations. A secondary axis was used to represent approximate nascent peptide length, assuming 3.5 Å per amino acid residue.

#### Polysome profiling

Polysome profiling was performed 24 hours after SNORA81 depletion to capture early translational effects in actively proliferating cells, before measurable proliferation or viability defects. Linear 5–50% sucrose gradients were prepared in buffer containing 20 mM Tris-HCl (pH 8.0), 140 mM KCl, 5 mM MgCl₂, 0.5 mM DTT, 0.1mg/mL cycloheximide and 0.5mg/mL heparin and were generated using a Gradient Master 108 (Biocomp) with program 81.5° 1.38minutes 18rpm. Gradients were stored at 4°C for no more than 24 h. For each condition, four 10cm dishes of OVCAR-3 cells were treated with 100ug/mL cycloheximide for 15 min at 37°C, washed with ice-cold PBS containing 100ug/mL cycloheximide, scraped, pooled and lysed in buffer containing 10 mM Tris-HCl (pH 7.5), 10 mM NaCl, 1.5 mM MgCl₂, 1.25% Triton X-100, 2.5% Tween-20, 0.5% sodium deoxycholate, 1X protease inhibitors (Roche), 1 mM PMSF, 40U RNase inhibitor (Université de Sherbrooke Protein Purification Platform) and 100ug/mL cycloheximide. Lysates were clarified by centrifugation, and equal protein amounts were loaded onto 5–50% sucrose gradients, which were centrifuged at 40,000 rpm for 3h at 4°C in an SW41 rotor. Gradients were fractionated with a Teledyne ISCO system and fractions corresponding to free, 40S, 60S, 80S and polysomal fractions (P1, two to three ribosomes; P2, four to six; P3, seven or more) were collected. A portion of each fraction was used for RNA extraction by ethanol precipitation with glycogen and potassium acetate, followed by on-column DNase treatment as described above, and 100 ng RNA per fraction was used for RNA-sequencing library preparation.

#### RNA sequencing

PolyA-selected RNA libraries were prepared from matched input samples generated both for the ribosome footprinting and polysome profiling experiments as well as from polysome fractions (40S, 80S, P1, P2 and P3). Libraries were prepared at the Université de Sherbrooke RNomics platform using the NEBNext^®^ Poly(A) mRNA Magnetic Isolation Module (E7490, NEB) and the NEBNext UltraExpress RNA Library Prep Kit (E3330, NEB) from 100ng total RNA per sample. Libraries were typically amplified using 10-12 PCR cycles. Polysome fraction libraries were amplified using 15 PCR cycles to compensate for lower input material. Libraries were sequenced on an Illumina’s NextSeq 500 using the NextSeq 500/500 High Output kit v2.5 (150 cycles; paired-end 2x75bp) or an Element AVITI System using the Cloudbreak Freestyle High Output 150 cycles kit 820-00022; 150 cycles; paired-end 2x75bp). polyA RNA sequencing was analysed as previously described^65^.

#### Transcript feature analysis

Protein-coding genes and their sequence were extracted from Ensembl’s human genome annotation and fasta files (hg38, v108) based on the biotype “protein_coding”. Briefly, one representative transcript was selected per gene based on the following criteria: 1) the transcript biotype is “protein_coding”; 2) the transcript annotation contains at least one annotated coding sequence (CDS), one annotated 5’ untranslated region (UTR) and one annotated 3’UTR; 3) in case of multiple transcripts sharing the previous criteria, transcript support level and nomenclature were used to choose one transcript (i.e., transcripts with the name ending with “-201” were favored over “-202”, and so on). In case of multiple CDSs annotated for a selected transcript, the longest CDS was selected as the representative CDS. Transcript, CDS, 5’ and 3’ UTR sequences were obtained using pybedtools^53^ sequence v0.10.0 with the -s option to force strandedness. The number of exons and introns per gene was also derived from the annotation file based on the selected transcript per gene.

#### G4 predictions

Potential RNA G4 within coding sequences were predicted using G4RNA screener webserver^49,50^. CDS sequences were analyzed using the default thresholds, and transcripts were classified as G4-positive only if at least one predicted G4 scored positive for all three metrics (cGcC, G4H and G4NN).

#### Gene ontology analysis

Gene ontology (GO) enrichment analysis was performed using the Molecular Signature Database (MSigDB) Gene Set Enrichment Analysis (GSEA) online platform^51,52^. Over-associated and under-associated STAR mRNAs were analyzed separately using the GO biological process (GO: BP) gene set collection. GO terms with a false discovery rate (FDR) q-values < 0.05 were considered significantly enriched. To reduce redundancy among enriched GO terms, rrvgo package in R^56^ was used based on GO term semantic relationships. GO term with a similarity threshold > 0.7 were grouped, and representative terms were selected based on enrichment significance. Redundant terms were collapsed into parent clusters and visualised. Enriched terms were subsequently assigned to manually curated cancer hallmark-related categories (Housekeeping and General, Development and Differentiation, Tumour Microenvironment and Immunity, Proliferation and Cell Cycle, Migration and Invasion, and Signalling and Gene Regulation). Cancer-associated genes were identified using CancerMine^47^.

#### Protein validation

Protein abundance was assessed 48 hours (TNF) and 72 hours (CDH1 and RBM25) after SNORA81 depletion to account for the slow turnover of ribosomes and downstream proteins. Western blotting was performed as described previously^13^. Cells were pelleted and lysed in buffer containing 150 mM NaCl, 50 mM Tris-HCl (pH 8.0), 0.5% NP-40, 0.5% sodium deoxycholate, 0.1% SDS, 10 mM sodium pyrophosphate and 50 mM EDTA supplemented with protease inhibitor cocktail (Roche), for 1h at 4 °C with rotation. Lysates were clarified by centrifugation, and protein concentration was determined by Bradford assays. Equal amounts of protein (20 µg) were denatured in Laemmli buffer, separated on 12% SDS–PAGE gels and transferred to Hybond-ECL membranes (GE Healthcare) at 100 V for 2h. Membranes were blocked overnight at 4°C in TBS-T containing 5% skimmed milk and incubated with primary antibodies diluted in blocking buffer (Anti-TNF, 1:1000, abcam ab9635; Anti-CDH1, 1:2000, BD Biosciences, catalogue 610182; Anti-RBM25, 1:2000, cell signaling 69882T; Anti-GAPDH, 1:2000, Novus Biologicals NB300-221) for 90 min at room temperature. Following washing, membranes were incubated with HRP-conjugated secondary antibodies (anti-rabbit, 1:2000, GE Healthcare, NA934; or anti-mouse, 1:2000, GE Healthcare, NA931) for 90 min at room temperature. Signals were detected using Clarity Western ECL substrate (Bio-Rad) and imaged on an LAS 4000 system (GE Healthcare).

### Quantification and statistical analysis

Statistical tests, biological replicate numbers and multiple-testing corrections are indicated in the corresponding Fig legends. For **Figure 3F**, exact p-values assessed by chi-square test between over and under groups for each fraction: 40S, P = 3.4 × 10⁻³²; 80S, P = 0.26; P1, P = 1.5 × 10⁻¹²; P2, P = 2.7 × 10⁻²¹; P3, P = 2.3 × 10⁻⁶. For **Figure 4A**, differences between distributions were assessed using two-sided Kolmogorov–Smirnov tests. P values comparing all with over and all with under: 5′ UTR, P = 0.013 and P = 0.025; CDS, P = 1.0 × 10⁻⁶ and P = 4.8 × 10⁻¹³; 3′ UTR, P < 1.0 × 10⁻¹⁶ and P = 4.3 × 10⁻¹².

