## Supplementary material for "A ribosomal pseudouridine drives an oncogenic translation program": Document S1

Faucher-Giguère et al 2026

Supplementary data

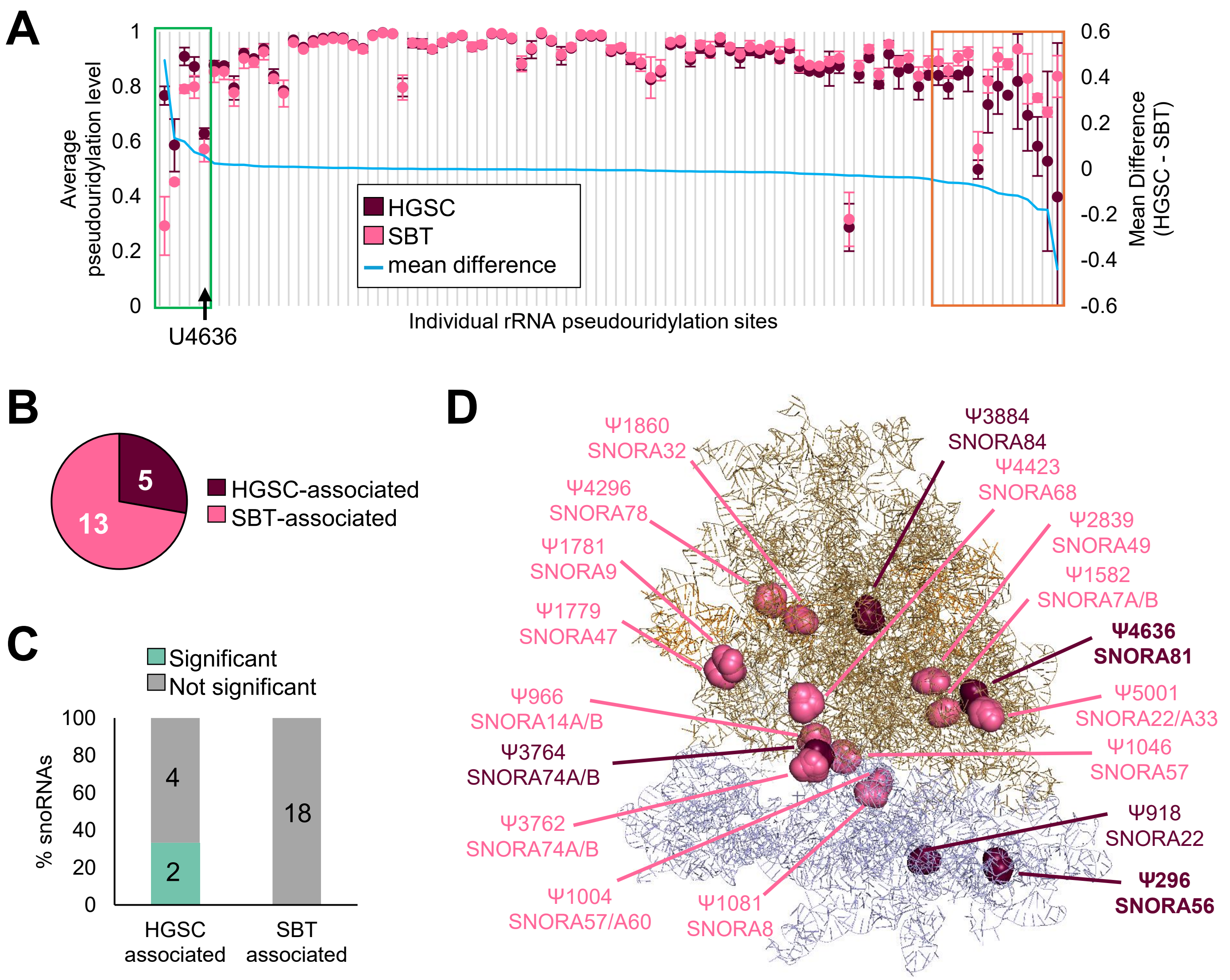


**Figure S1 (related to Figure 1). A subset of rRNA pseudouridylation sites and their guide snoRNAs differ between HGSC and SBT.**

**(A)** Pseudouridylation at individual rRNA sites in five ovarian tissue samples, comprising two high-grade serous carcinomas (HGSC) and three serous borderline tumours (SBT). Points show the mean modified fraction ± s.d. for HGSC (dark magenta) and SBT (light pink). The blue line shows the difference between group means (HGSC minus SBT). Boxes indicate sites with a difference in mean modified fraction greater than 0.05 (green) or lower than -0.05 (orange). U4636 is indicated. **(B)** Distribution of the 18 sites identified in a, comprising five HGSC-associated and thirteen SBT-associated sites. **(C)** Differential abundance of guide snoRNAs corresponding to the sites identified in b, based on previously published RNA-sequencing data (Faucher-Giguère et al. 2022). Bars show the percentage of guide snoRNAs that were significantly differentially abundant between HGSC and SBT (green) or not significantly different (grey); numbers indicate snoRNA counts. Multiple snoRNAs can guide the same rRNA site, so the number of guide snoRNAs differs from the number of sites in b. Significance was determined using DESeq2 (|log2 fold change| > 0.5, adjusted $P < 0.05$). **(D)** Structural localization of HGSC-associated (dark magenta) and SBT-associated (light pink) pseudouridylation sites on the human ribosome. Bold labels identify sites for which at least one corresponding guide snoRNA was significantly differentially abundant between HGSC and SBT, as defined in c.

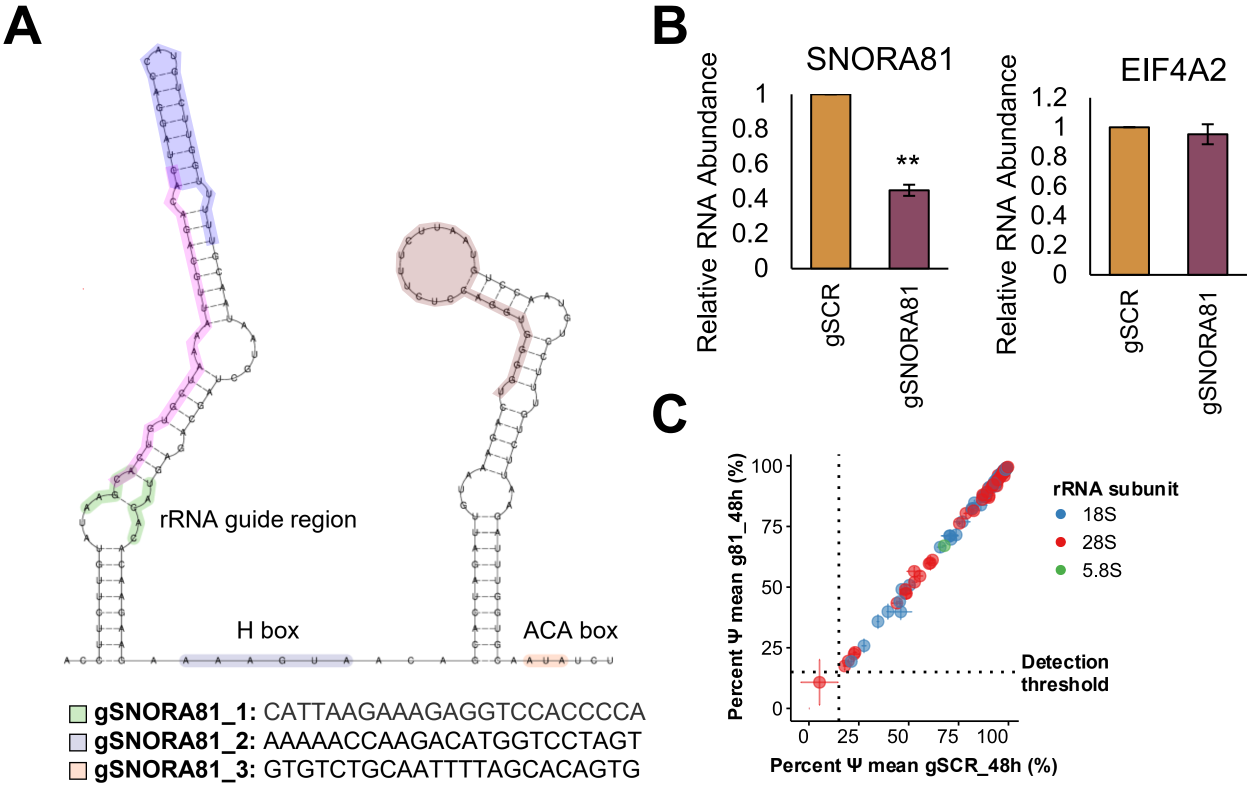


**Figure S2 (related to Figure 1). Validation of CRISPR–Cas13d-mediated SNORA81 depletion.**

**(A)** Predicted SNORA81 secondary structure (extracted from SNODB) showing the rRNA guide region, H box, ACA box and the three CRISPR–Cas13d guide RNA target sites (Guides 1–3); guide sequences (gSNORA81_1–3) are listed below. **(B)** RT–qPCR validation of CRISPR–Cas13d-mediated SNORA81 depletion (left) and unchanged RNA abundance of the host gene EIF4A2 (right) in gSNORA81 versus gSCR cells, shown as relative RNA abundance. Data are mean ± s.d. of n = 3 biological replicates; significance by two-sided T-test; SNORA81, P = 0.0011; EIF4A2, not significant (P > 0.05). **(C)** Global rRNA pseudouridylation after SNORA81 depletion. Each point is the mean Ψ level (%) at one site in gSNORA81 (g81_48h, y axis) versus gSCR (gSCR_48h, x axis), coloured by rRNA subunit (18S, blue; 28S, red; 5.8S, green); error bars are s.d. across n = 3 biological replicates. Dotted lines mark the detection threshold.

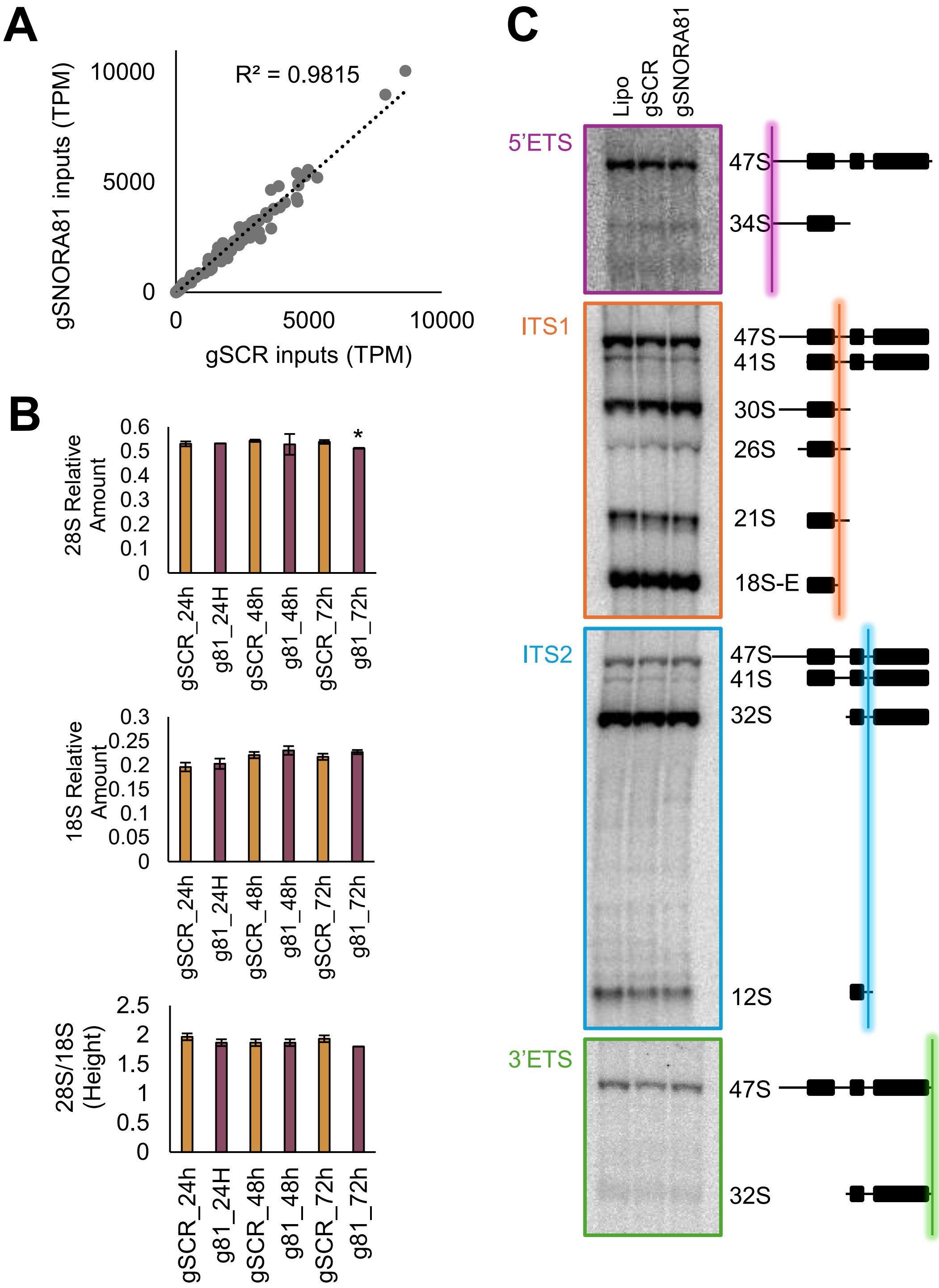


**Figure S3 (related to Figure 1). SNORA81 depletion alters translation without disrupting ribosome production.**

**(A)** Scatter plot of ribosomal-protein mRNA abundance in input RNA from gSNORA81 versus gSCR cells. Each point represents one ribosomal-protein gene. Abundance was measured by RNA-seq and is expressed as transcripts per million (TPM). The dotted line shows the linear regression, with $R^2$ indicated. **(B)** Relative abundance of mature 28S and 18S rRNA and the 28S/18S ratio at the indicated times after SNORA81 depletion. Data are mean ± s.d. from n = 3 biologically independent samples. Significance was assessed using two-sided t-tests. The abundance of 28S rRNA differed significantly between gSCR and gSNORA81 cells at 72 h (P = 0.02). **(C)** Northern blot analysis of pre-rRNA processing intermediates detected using probes against the 5′ ETS, ITS1, ITS2 and 3′ ETS regions after SNORA81 depletion.

**A**

| Structure | Cell type | Resolution (Å) | Ψ4636 state |
|---|---|---|---|
| 8QOI | HeLa | 1.9 | Pseudouridylated and confirmed by MS |
| 9PBE | 293A | 2.2 | Pseudouridylated |
| 9PKG | 293A | 3.3 | Pseudouridylated |
| 8QYX | HeLa | 1.78 | unmodified |
| 9SPF | Human induced pluripotent stem cells | 2.4 | unmodified |
| 7F5S | HEK293T (KO METTL18) | 2.7 | unmodified |
| 9BDN | Rabbit reticulocyte lysate (RRL) | 3.1 | unmodified |
| 6SGC | Rabbit reticulocyte lysate (RRL) | 2.8 | unmodified |

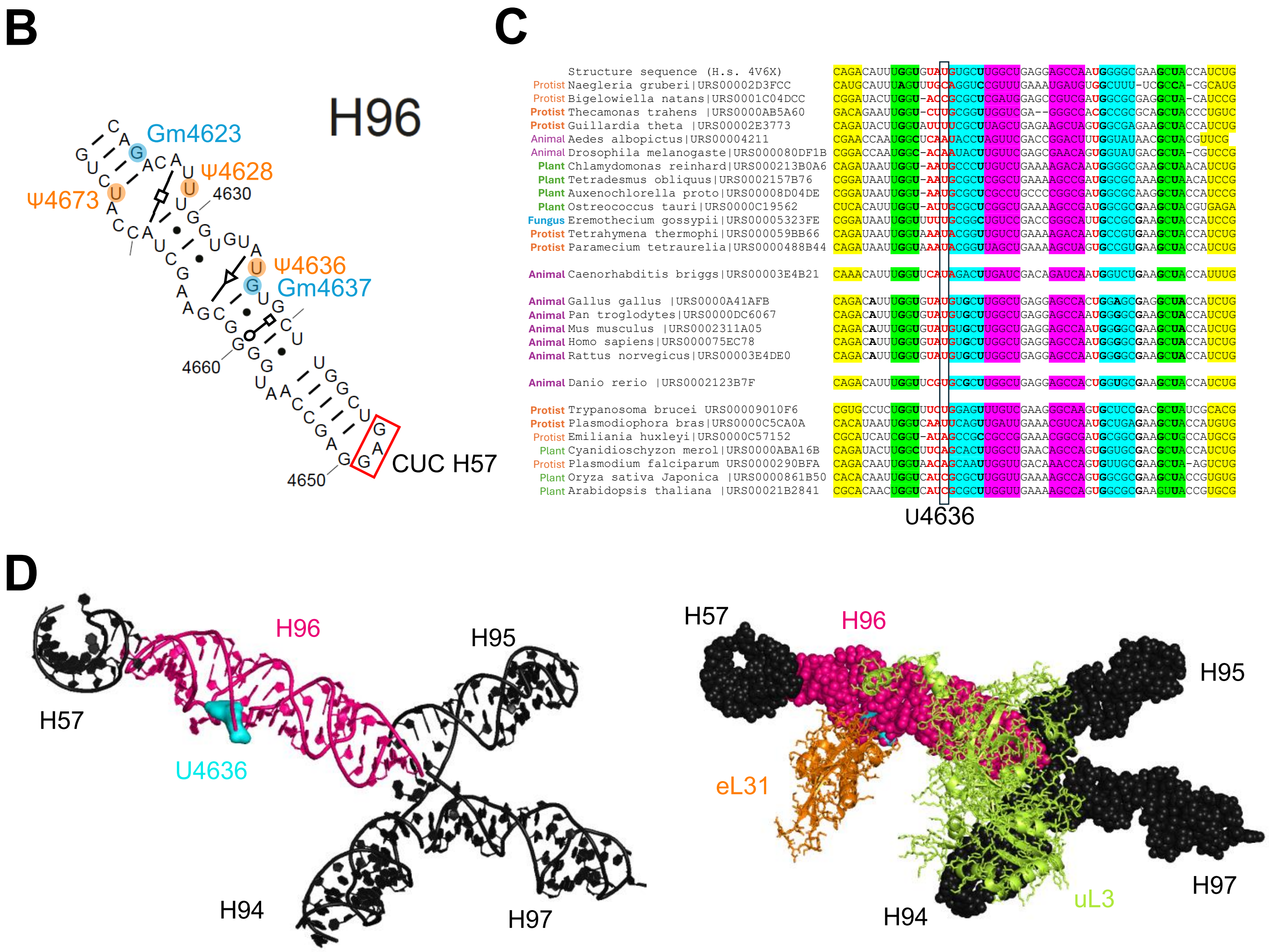


**Figure S4 (related to Figure 1). Conservation and structural organization of the H96 region containing U4636.**
**(A)** Summary of available mammalian ribosome structures analyzed to assess the molecular environment surrounding U4636. **(B)** Secondary-structure representation of H96 showing pseudouridylation residues in orange and 2'-O-methylation in blue. The nucleotide involved in the kissing-loop interaction with H57 is indicated by a red box. **(C)** Sequence alignment of the H96 region across representative eukaryotes. U4636 is conserved in most mammalian sequences examined but shows greater variation across other eukaryotic lineages (Ribovision2, McCann, 2024). **(D)** Three-dimensional representation of the H94–H97 four-way junction containing H96, shown as RNA alone (left) and with eL31 and uL3 (right), highlighting the position of H96 within the larger rRNA structural framework.

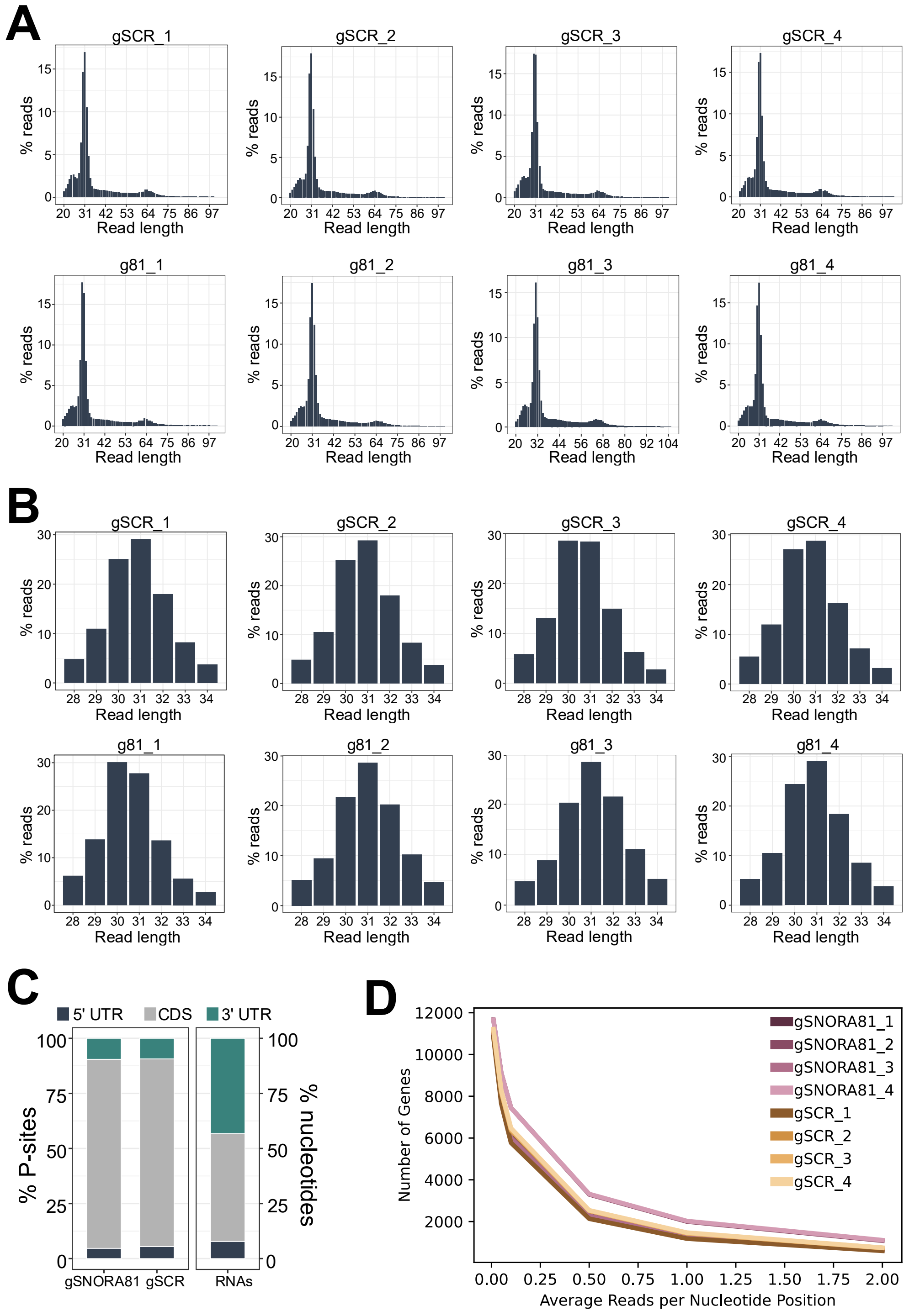


**Figure S5 (related to Figure 2). Quality-control analysis of ribosome profiling libraries following SNORA81 depletion.**

**(A)** Read-length distribution of all ribosome-protected fragments (RPFs) in four control (gSCR_1-4) and four SNORA81-depleted (gSNORA81_1-4) biological replicates. Values are shown as the percentage of reads at each length. **(B)** Read-length distribution after filtering to the canonical RPF size range retained for downstream P-site assignment and pausing analyses (28-34 nt), shown for the same replicates. **(C)** Distribution of P-sites across transcript regions (5′ UTR, CDS and 3′ UTR) in gSCR and gSNORA81 samples, showing predominant coding-sequence enrichment. The corresponding distribution of total RNA-seq reads is shown for comparison. **(D)** Gene-coverage analysis showing the number of genes retained across increasing mean per-gene CDS coverage thresholds for each ribosome-profiling replicate.

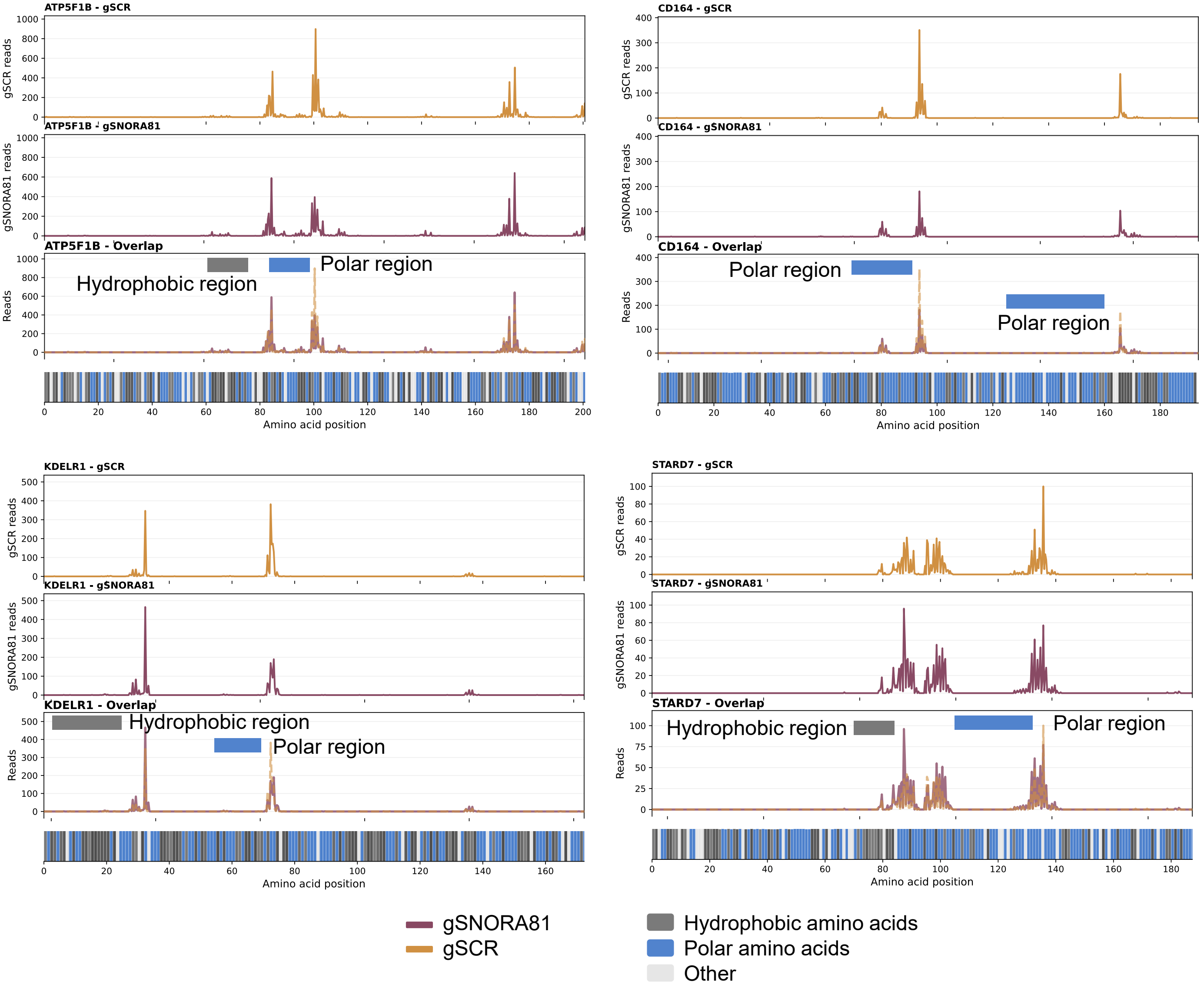


**Figure S6 (related to Figure 2) Individual gene coverage plots showing pause sites and amino acid properties.**

Each panel shows ribosome profiling coverage and amino acid properties for genes with differential pause sites enriched in specific tunnel environments. Coverage tracks are shown for gSCR (yellow), SNORA81 (purple) and overlay of both conditions. The amino acid track uses a gradient color scheme: grey (hydrophobic, where darker is more hydrophobic) and blue (polar), with light grey for other residues. The x-axis indicates amino acid position.

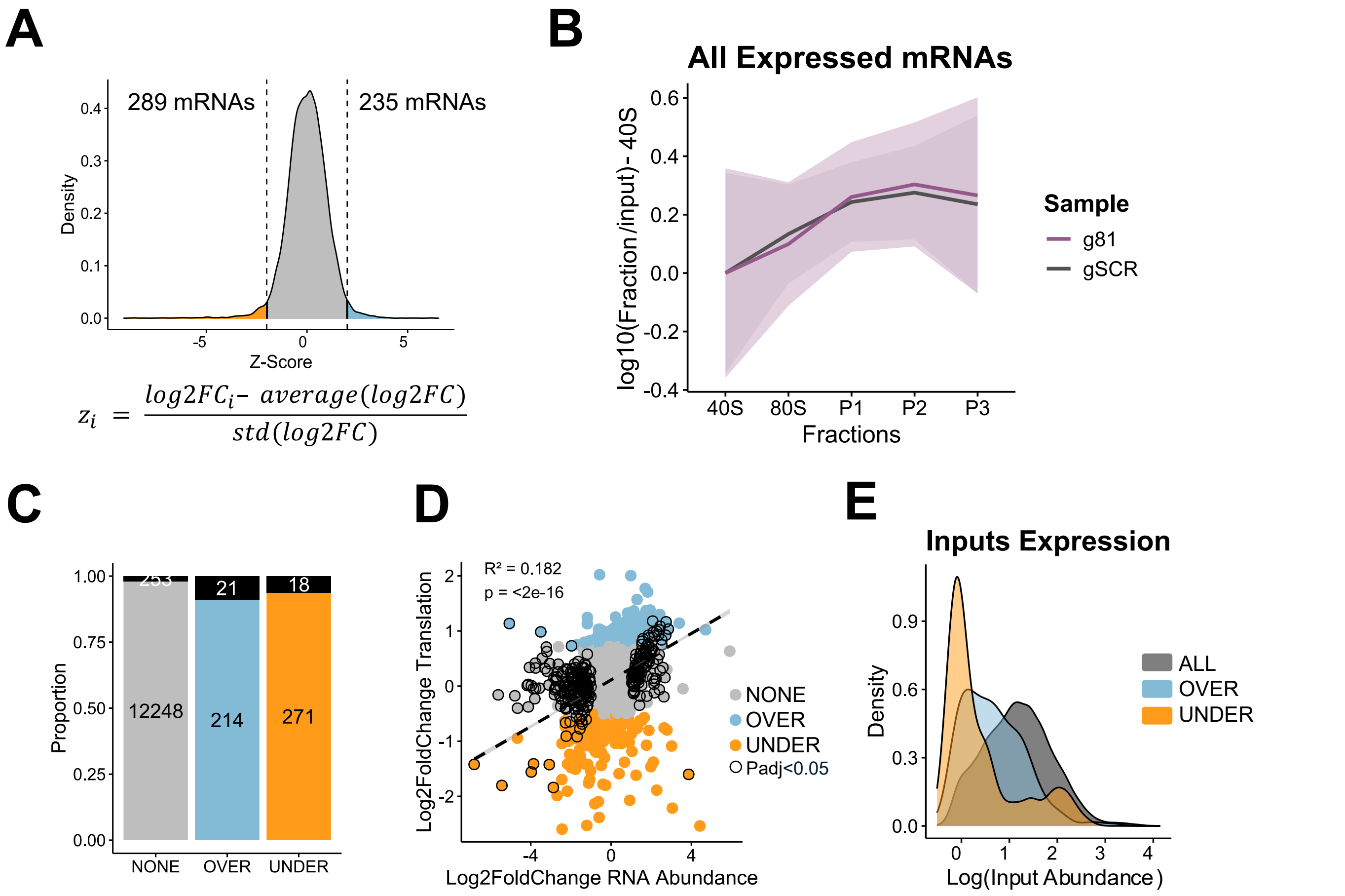


**Figure S7 (related to Figure 3) SNORA81 depletion alters ribosome association without broad changes in mRNA abundance.**
**(A)** Distribution of translation z-scores after SNORA81 depletion, calculated from the average log2 fold change in ribosome association between gSNORA81 and gSCR samples. Dashed lines indicate the thresholds used to define mRNAs with decreased ($z \leq -2$; 289 mRNAs, orange) or increased ($z \geq 2$; 235 mRNAs, blue) ribosome association. **(B)** Global mRNA distribution across polysome fractions after SNORA81 depletion. Lines show mean log10(fraction/input) values across 40S, 80S and polysome fractions for gSCR (grey) and gSNORA81 (purple) samples; shaded regions indicate s.d. **(C)** Proportion of mRNAs classified as unchanged, over-associated or under-associated by polysome profiling. Black segments indicate mRNAs that are also differentially abundant in input RNA-seq data, defined as adjusted $P < 0.05$ and absolute log2 fold change > 0.5. **(D)** Scatter plot comparing log2 fold change in total mRNA abundance (input RNA-seq; x axis) with log2 fold change in ribosome association (y axis). Points indicate unchanged mRNAs (grey), over-associated mRNAs (blue), under-associated mRNAs (orange) and differentially abundant mRNAs (black); the correlation coefficient and P value are shown. **(E)** Distribution of input mRNA abundance, shown as log10 (input abundance), for all detected mRNAs (grey), over-associated mRNAs (blue) and under-associated mRNAs (orange).

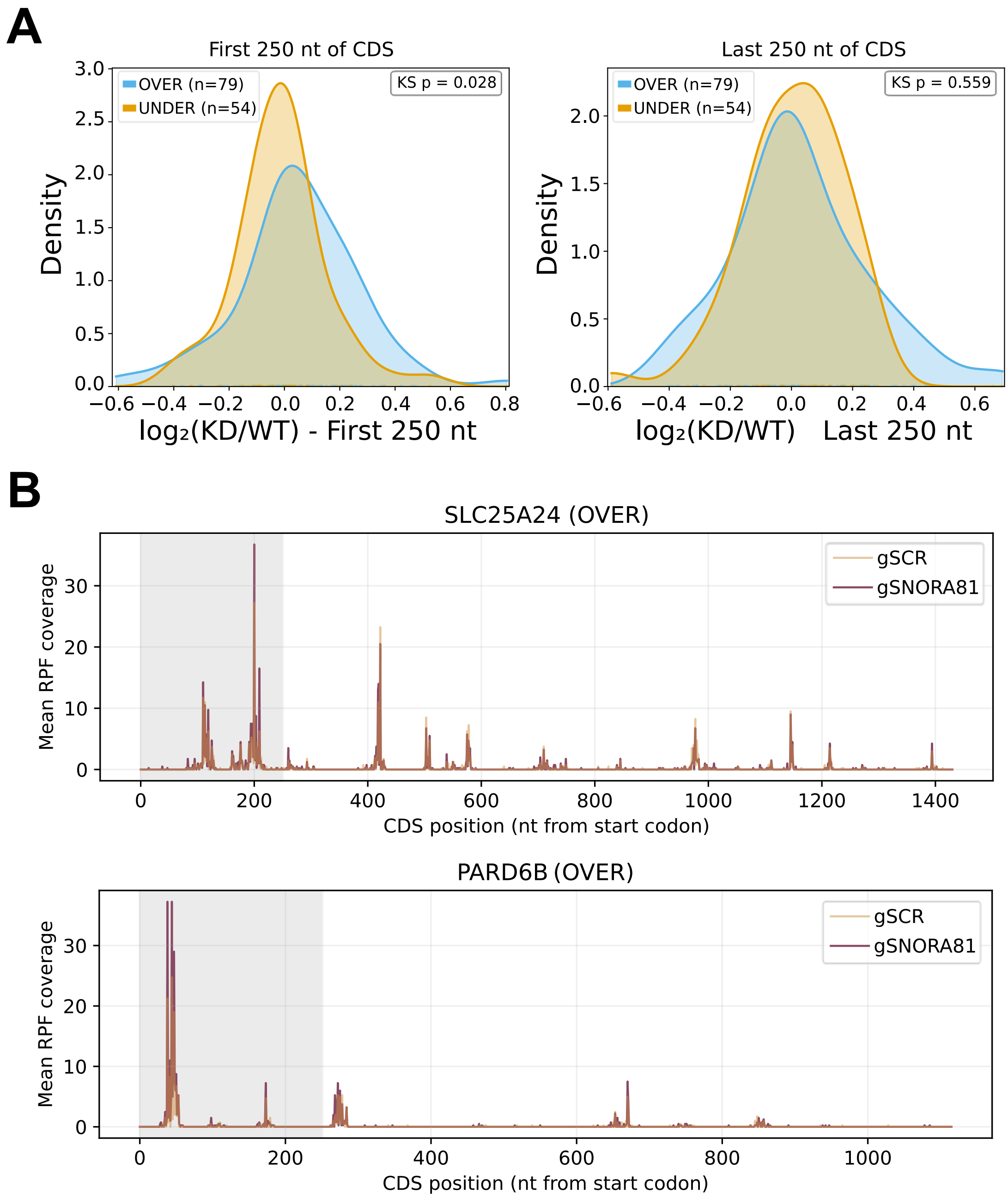


**Figure S8 (related to Figure 3). SNORA81 depletion differentially affects ribosome occupancy near coding-region boundaries**.

**(A)** Density distributions of the log2 ratio of normalized ribosome-footprint reads in SNORA81-depleted versus control cells within the first 250 nt (left) and last 250 nt (right) of coding sequences (CDS). Over STAR mRNAs are shown in blue and under STAR mRNAs in orange. For each transcript, ribosome protected fragments (RPF) counts within each window were normalized to total CDS ribosome-footprint counts. Analyses were performed with a minimum CDS coverage threshold of 0.1. Kolmogorov–Smirnov (KS) tests were used, P value are indicated in the plots. **(B)** Metagene analysis of OVER STAR mRNAs showing mean normalized RPF coverage across coding sequences in SNORA81-depleted and control (gSCR). The x-axis indicates position relative to start codon, and the y-axis indicates mean normalized RPF coverage.

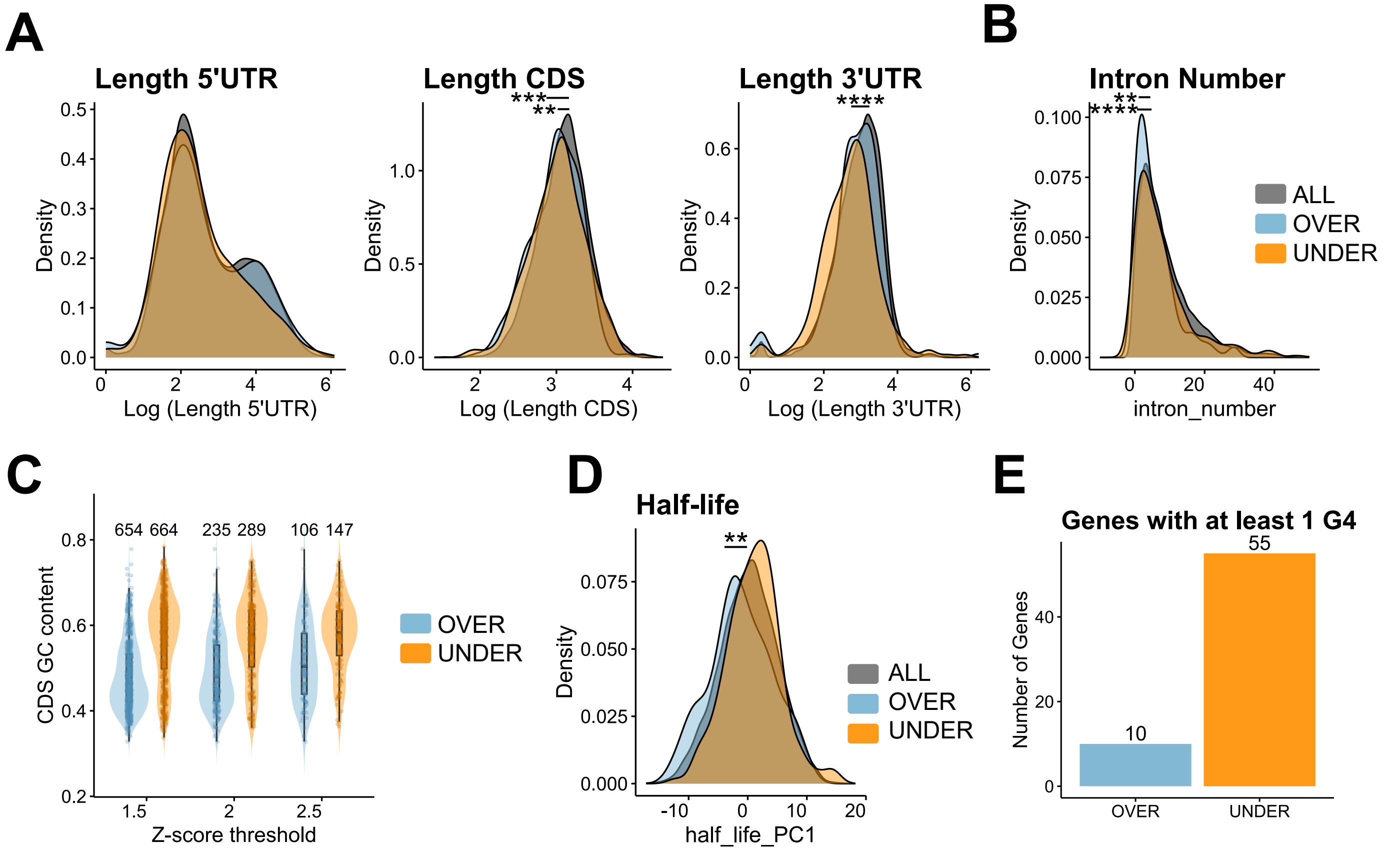


**Figure S9 (related to Figure 4). Transcript features associated with SNORA81-dependent ribosome association.**

**(A)** Density plots showing the log-transformed lengths of the 5′ UTR, coding sequence (CDS) and 3′ UTR for all detected mRNAs (grey), over-associated mRNAs (blue) and under-associated mRNAs (orange) after SNORA81 depletion. **(B)** Density plot showing intron number for all detected mRNAs (grey), over-associated mRNAs (blue) and under-associated mRNAs (orange). **(C)** Robustness of CDS GC-content association across alternative STAR mRNA Z-score thresholds. Violin plot showing CDS GC content distributions for over (blue) and under (orange) associated mRNAs classified using alternative z-score threshold (|Z-score| ≥ 1.5, 2 and 2.5). Numbers above each distribution indicate the number of transcripts included in each group. **(D)** Density plot showing mRNA half-life, represented as principal component analysis derived PC1 value (Agarwal, et al. 2022), for all detected mRNAs (grey), over-associated mRNAs (blue) and under-associated mRNAs (orange). **(E)** Number of over-associated and under-associated mRNAs containing at least one predicted G-quadruplex (G4) within the CDS. G4s were predicted using G4RNA Screener (Garant et al., 2017), and transcripts were classified as G4-positive only if they scored positive by all three metrics: cGcG, G4H and G4NN. Statistical significance in **A, B, D** was assessed using two-sided Kolmogorov–Smirnov tests; Exact P values were: a, CDS, ALL vs OVER ($P = 5.41 \times 10^{-4}$) and ALL vs UNDER ($P = 1.68 \times 10^{-3}$); 3′ UTR, ALL vs UNDER ($P = 2.27 \times 10^{-9}$). b, intron number, ALL vs OVER ($P = 2.92 \times 10^{-6}$) and ALL vs UNDER ($P = 8.36 \times 10^{-3}$). c, mRNA half-life, ALL vs OVER ($P = 3.43 \times 10^{-3}$).

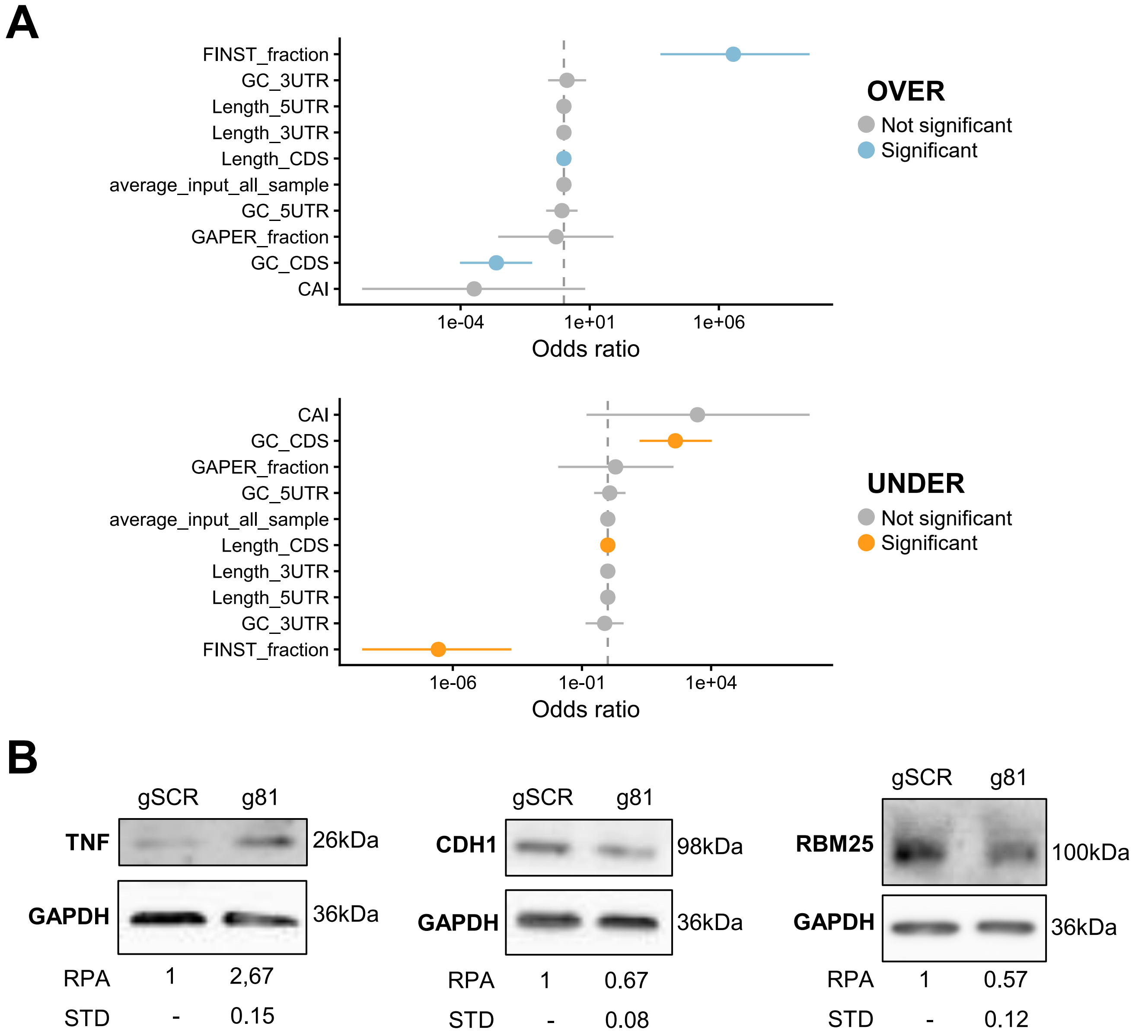


**Figure S10 (related to Figure 4 and 5). Features associated with STAR mRNAs and validation of selected protein changes following SNORA81 depletion.**
**(A)** Forest plot showing odds ratios from multivariate logistic regression comparing over (top) and under (bottom) STAR transcripts versus all expressed mRNAs (n = 13,025 genes). The model included GC content (5'UTR, 3'UTR, CDS), transcript lengths (5'UTR, 3'UTR, CDS), mRNA abundance (average from input), codon adaptation index (CAI), and codon signature metrics (FINST and GAPER fractions). Points represent odds ratios, with error bars indicating 95% confidence intervals Odds ratios > 1 (blue) indicate features associated with over STAR mRNAs, while odds ratios < 1 (orange) indicate features associated with under associated STAR mRNAs. **(B)** Representative blot showing the effect of SNORA81 depletion on protein abundance of selected STAR mRNAs. TNF, CDH1 and RBM25 protein levels were determined before and after SNORA81 knockdown. GAPDH was used as loading control. Average relative protein levels from three biological replicates are indicated below the representative blot and estimated molecular weight are indicated on the side.

**Table S1: cas13d knockdown guide sequences**

| guide | sequence |
|---|---|
| gSCR | ACAGGATGTGTAGCACTGATGCGA |
| gSNORA81_1 | CATTAAGAAAGAGGTCCACCCCA |
| gSNORA81_2 | AAAAACCAAGACATGGTCCTAGT |
| gSNORA81_3 | GTGTCTGCAATTTTAGCACAGTG |

**Table S2: Primers used in this study for RT-qPCR**

| Primers | Function | Forward sequences | Reverse sequences |
|---|---|---|---|
| MRPL19 | Housekeeping | AAGGAGAAAAGTACTCCACATTCCAGAG | TGGGTCAGCTGTAGTAACACGA |
| RPPH1 | Housekeeping | TGAGTGCGTCCTGTCACTCCA | ACTCACCTCCCCGAAGCTCA |
| YWHAZ | Housekeeping | TCCCCAATGCTTCACAAGCAGA | TCTTGTCATCACCAGCGGCAA |
| SNORA81 | snoRNA | GTGCTAAAATTGCAGACACTAGGACCA | GCACCAAATCTTAAGACAAAGGACATTGGA |
| EIF4A2 | Host gene | GTACACCCGGGAGAGTGTTT | ACGGCTCAACATTTCATCTG |
